# A cross-scale analysis to understand and quantify effects of photosynthetic enhancement on crop growth and yield

**DOI:** 10.1101/2022.07.06.498957

**Authors:** Alex Wu, Jason Brider, Florian A. Busch, Min Chen, Karine Chenu, Victoria C. Clarke, Brian Collins, Maria Ermakova, John R. Evans, Graham D. Farquhar, Britta Forster, Robert T. Furbank, Michael Gorszmann, Miguel A. Hernandez, Benedict M. Long, Greg Mclean, Andries Potgieter, G. Dean Price, Robert E. Sharwood, Michael Stower, Erik van Oosterom, Susanne von Caemmerer, Spencer M. Whitney, Graeme L. Hammer

**Affiliations:** ARC Centre of Excellence for Translational Photosynthesis, Centre for Crop Science, Queensland Alliance for Agriculture and Food Innovation, The University of Queensland, Brisbane, Queensland, Australia; ARC Centre of Excellence for Translational Photosynthesis, Division of Plant Science, Research School of Biology, The Australian National University, Acton, Australian Capital Territory, 2601, Australia; School of Biosciences, University of Birmingham, Birmingham, B15 2TT, UK; Birmingham Institute of Forest Research, University of Birmingham, Birmingham, B15 2TT, UK; ARC Centre of Excellence for Translational Photosynthesis, School of Life and Environmental Science, Faculty of Science, University of Sydney, Sydney NSW2006, Australia; Hawkesbury Institute for the Environment, Western Sydney University, Richmond, NSW, 2154, Australia

**Author notes:** **Author contributions:** AW, GH, GF, and SvC designed the study. AW and GH wrote the manuscript. AW updated and tested the model; AW, JB, GM and MS updated and tested the software. AW and JB constructed the simulation files and performed all model simulations with input from GH, KC, BC, GM, AP, and EvO. FB, MC, TC, ME, JE, GF, BF, RF, MG, MH, BL, DP, RS, SvC, and SW contributed detailed photosynthesis knowledge justifying photosynthetic parameter changes used in the simulation files. All authors provided feedback and edited the paper.

**Keywords:** crop growth modelling, crop production, yield improvement, APSIM, electron transport-limited photosynthesis, enzyme-limited photosynthesis

## Abstract

Photosynthetic manipulation provides new opportunities for enhancing crop yield. However, understanding and quantifying effectively how the seasonal growth and yield dynamics of target crops might be affected over a wide range of environments is limited. Using a state-of-the-art cross-scale model we predicted crop-level impacts of a broad list of promising photosynthesis manipulation strategies for C_3_ wheat and C_4_ sorghum. The manipulation targets have varying effects on the enzyme-limited (*A*_c_) and electron transport-limited (*A*_j_) rates of photosynthesis. In the top decile of seasonal outcomes, yield gains with the list of manipulations were predicted to be modest, ranging between 0 and 8%, depending on the crop type and manipulation. To achieve the higher yield gains, large increases in both *A*_c_ and *A*_j_ are needed. This could likely be achieved by stacking Rubisco function and electron transport chain enhancements or installing a full CO_2_ concentrating system. However, photosynthetic enhancement influences the timing and severity of water and nitrogen stress on the crop, confounding yield outcomes. Strategies enhancing *A*_c_ alone offers more consistent but smaller yield gains across environments, *A*_j_ enhancement alone offers higher gains but is undesirable in less favourable environments. Understanding and quantifying complex cross-scale interactions between photosynthesis and crop yield will challenge and stimulate photosynthesis and crop research.

**Summary Statement:** Leaf–canopy–crop prediction using a state-of-the-art cross-scale model improves understanding of how photosynthetic manipulation alters wheat and sorghum growth and yield dynamics. This generates novel insights for quantifying impacts of photosynthetic enhancement on crop yield across environments.

## Introduction

New strategies to improve grain yield in globally important staple crops are needed urgently if production is to keep pace with growing demand (Fischer *et al*., 2014, Ray *et al*., 2013). Improving crop resource use efficiencies and crop growth rates are promising avenues and photosynthesis has emerged as one of the major traits of interest (Evans, 2013, Hammer *et al*., 2020, Long *et al*., 2015, von Caemmerer & Furbank, 2016). Manipulation of a number of key proteins involved has been achieved in transgenic plants with causal enhancement in leaf CO_2_ assimilation rates (Ermakova *et al*., 2019, Salesse-Smith *et al*., 2018). Researchers have also modelled consequences of leaf CO_2_ assimilation rate with enhanced Rubisco and installation of a cyanobacterial CO_2_ concentrating system (Price *et al*., 2011, Sharwood *et al*., 2016b). Numerous studies reported large changes in plant-level attributes with enhanced photosynthesis (Simkin *et al*., 2019). Such promising results have been used in projecting large gains in crop production, however, more understanding and quantification of seasonal crop growth and yield dynamics are needed for assessing yield impacts credibly.

This requires understanding of interactions between perturbed leaf photosynthetic and crop growth rates, crop developmental processes, crop resources supply and demand, regulation of leaf photosynthesis by status of the crop, and environment context dependencies (Hammer *et al*., 2010, Wu *et al*., 2016). Free-air CO_2_ enrichment studies provide indirect evidence of potential C_3_ crop yield improvement with enhanced photosynthesis from elevated CO_2_ under non-stress conditions (Ainsworth & Long, 2021), but causal physiological links between leaf photosynthesis and crop yield need to be further unravelled and assessed in broader environmental conditions to predict yield credibly (Fischer *et al*., 1998). Interactions between the growing crop in contrasting environments can generate complex crop growth and yield consequences (Hammer *et al*., 2016). Ideally, photosynthetically enhanced plants need to be tested using multi-environment trials (i.e. field testing of target crops at several representative production locations over several years) for better understanding and quantification of growth and yield dynamics, but such an approach is mostly inaccessible. Absence of such information hampers efforts to maximize yield improvement (Fischer *et al*., 2014).

Crop growth modelling is a useful method for predicting the growth and yield dynamics and their interactions with the growing environment. Recent research thrusts in crop growth modelling paved the way for achieving the necessary leaf-to-crop connection (Chew *et al*., 2017, Hammer *et al*., 2019, Marshall-Colon *et al*., 2017, Wu *et al*., 2016). Models that incorporate complexities associated with interactions between leaf photosynthetic rates, diurnally changing temperature, solar radiation, and with-in canopy light environment can be used to predict daily canopy photosynthetic and/or crop growth rates (e.g. de Pury & Farquhar (1997); Hammer & Wright (1994); Song *et al*. (2013); Wu *et al*., (2018)). Crop models that incorporate complex interactions between crop phenology, canopy development, growth, and effects of whole-crop water and nitrogen supply/demand on growth and development processes (Brown *et al*., 2014, Hammer *et al*., 2010) provide information for predicting leaf and canopy photosynthesis over the crop life cycle. A previous study using a rice crop model combined with a photosynthesis model suggested that photosynthetic manipulation can generate moderate to large biomass gains under well-watered conditions, but even larger gains with water limitation (Yin & Struik, 2017). However, the results for water-limited situations contrast with known interactions between enhanced crop growth and water limitation (Hammer *et al*., 2010). There is a need to better understand and quantify impacts of photosynthetic bioengineering strategies on seasonal crop growth and yield dynamics in a wide range of environments. Models that have been demonstrated to predict field crop data in a wide range of environments by capturing the two-way interactions between leaf photosynthesis and crop growth and yield processes will be needed (Wu *et al*., 2019, Wu *et al*., 2016).

This study aims to understand and quantify dynamics of wheat and sorghum growth and yield with enhanced photosynthesis using a state-of-the-art cross-scale crop growth model (Wu *et al*., 2019). The three objectives are: (1) compose a list of promising photosynthetic enhancement strategies for C_3_ and C_4_ photosynthesis and model their effects on the enzyme-limited (*A*_c_) and electron transport-limited (*<u>A</u>*_j_) rates of CO_2_ assimilation using a generic photosynthesis model applicable for C_3_, C_4_ and single-cell CCM pathways; (2) conduct detailed leaf-to-yield analysis to understand consequences of perturbed leaf photosynthesis on crop growth and yield dynamics throughout the crop life cycle over a broad range of crop production environments; (3) conduct a case study on yield impacts for Australian wheat and sorghum. Manipulation targets promising the greatest gains in crop production are identified and implications of this modelling work for photosynthesis research and crop improvement decision-making are discussed.

## Materials and Methods

### Cross-scale modelling overview

This study aims to understand and quantify growth and yield dynamics of wheat and sorghum crops with enhanced photosynthesis in real production environments. The cross-scale model (Wu *et al*., 2019) used in this study allows two-way connections between the key biochemical processes of photosynthesis, which include the enzyme-limited (*A*_c_) and electron transport-limited (*<u>A</u>*_j_) rates of CO_2_ assimilation (Farquhar *et al*., 1980, von Caemmerer, 2000), a sun– shade canopy photosynthesis model (Wu *et al*., 2018), and the APSIM crop growth models (Brown *et al*., 2014, Hammer *et al*., 2010). The well parameterised APSIM models capture physiological determinants of crop growth, development and yield process and their interactions with the environments well (Brown *et al*., 2014, Hammer *et al*., 2010). With the cross-scale modelling framework advances, it has been demonstrated to predict photosynthesis and field crop yield in a wide range of environments (Wu *et al*., 2019).

At the leaf level, an expanded generic leaf photosynthesis model coupled with CO_2_ diffusion processes and leaf energy balance (Wu *et al*., 2019) was used to model the C_3_, C_4_ and the cyanobacterial CCM pathways. For modelling the CCM pathway, it captures the active transport of dissolved inorganic carbon into the mesophyll and its take-up by specialized protein micro-compartments, carboxysomes, that concentrate CO_2_ around the encapsulated Rubisco (Price *et al*., 2013, Rae Benjamin *et al*., 2013). The photosynthesis–CO_2_ diffusion model has the capacity to simulate effects of varying light, temperature, leaf nitrogen content, and transpiration on leaf CO_2_ assimilation rate (Wu *et al*., 2019). All photosynthesis–CO_2_ diffusion model equations are given in Appendix A and baseline parameter values are given in Table S5.

Key photosynthetic manipulation targets are detailed in a following section and approaches to model them using the photosynthesis–CO_2_ diffusion and canopy models are given in Table 1. Some approaches are common for both plant types, while some are plant type specific (e.g. installation of the cyanobacterial CCM in C_3_ wheat, but not C_4_ sorghum). Simulations in this work include leaf-level photosynthetic response to intercellular CO_2_ (*A–C*_i_), diurnal canopy photosynthesis and biomass accumulation, whole-crop growth, development, and yield dynamics (from sowing of the crop to harvest) in a broad range of production environments (Table S4).

**Table 1.** Comprehensive information on modelling effects of photosynthetic manipulation at the leaf level. This includes key aspects of leaf photosynthetic function in C_3_ wheat and C_4_ sorghum, the associated manipulation outcomes, bioengineering strategies, and representation using the generic leaf photosynthesis model coupled with CO2 diffusion processes (Appendix A). Component(s) of the photosynthetic machinery relevant to the bioengineering strategies are indicated in the schematic diagram of photosynthetic pathways (Figure 1). The relative changes in the photosynthetic parameters are applied to the baseline set (Table S5).

| Key aspects of leaf photosynthetic function | Manipulation outcome | Bioengineering strategy | Modelling of bioengineering targets with the generic leaf photosynthesis model |
| --- | --- | --- | --- |
| Rubisco function | (1.1) Rubisco with desirable C<sub>4</sub> NADP-ME maize catalytic properties (in C<sub>3</sub> wheat) | Engineering the endogenous Rubisco to achieve increased $k_{cat}^c$ and carboxylation efficiency (CE: $k_{cat}^c/[K_c(1 + O/K_o)]$) as found in C<sub>4</sub> maize Rubisco. | +50% $k_{cat}^c$, but only a +30% CE (as a result of concomitant changes in $K_c$ (+18%) and $K_o$ (+3%)) (Sharwood *et al.*, 2016a, Sharwood *et al.*, 2016b); assumed no change in $S_{c/o}$ or temperature response of any of the Rubisco catalytic properties; $k_{cat}^c$ increase is modelled by a proportional 50% increase in $V_{cmax}$ assuming the amount of Rubisco enzyme is the same and no change in the activation state. |
| | (1.2) Reducing wasteful photorespiration with higher specificity for CO<sub>2</sub> over O<sub>2</sub> ($S_{c/o}$) (in C<sub>3</sub> wheat) | Engineering the endogenous Rubisco to increase $S_{c/o}$. | +20% $S_{c/o}$ (Martin-Avila *et al.*, 2020); assumed no change in $V_{cmax}$, $K_c$ and $K_o$ or temperature response or any of the Rubisco catalytic properties. |
| | (1.3) Increased Rubisco content (in C<sub>4</sub> sorghum) | Overexpressing the endogenous Rubisco. | +20% $V_{cmax}$ (Salesse-Smith *et al.*, 2018). |
| | (1.4) Reduced affinity of Rubisco for oxygen (in C<sub>4</sub> sorghum) | Engineering the endogenous Rubisco to increase $K_o$ (von Caemmerer & Furbank, 2016). | +20% increase in $K_o$; concomitant change in $S_{c/o}$; assumed no change in $V_{cmax}$ and $K_c$ or temperature response of any of the Rubisco catalytic properties. |
| | (2) “Better” Rubisco:<br>• C<sub>3</sub>: Rubisco having C<sub>4</sub> NADP-ME maize like catalytic properties and higher $S_{c/o}$ | C<sub>3</sub> wheat:<br>Stacking manipulation in $k_{cat}^c$, CE and $S_{c/o}$ from combining Targets 1.1 and 1.2. | $k_{cat}^c$ (or $V_{cmax}$), CE, and $S_{c/o}$ modified as described for Targets 1.1 and 1.2. |
| | | C<sub>4</sub> sorghum: | $V_{cmax}$ and $K_o$ (affecting $S_{c/o}$) modified as described for Targets 1.3 and 1.4. |
| | • C<sub>4</sub>: Increased amount and improved catalytic properties | Stacking manipulation in Rubisco amount and $S_{c/o}$ from combining Targets 1.3 and 1.4. | |
| CO<sub>2</sub> delivery | (3) Improved diffusion of CO<sub>2</sub> into the mesophyll | Altering membrane permeability to CO<sub>2</sub> by adding CO<sub>2</sub> permeable plasma membrane intrinsic proteins, aquaporins to increase mesophyll conductance (Groszmann *et al.*, 2017). | +20% mesophyll conductance ($g_m$) (Groszmann *et al.*, 2017); assumed no change to the temperature response of $g_m$. |
|  | (4.1) Cyanobacterial CCM: active transport of dissolved inorganic carbon to reduce the drawdown of CO<sub>2</sub> in the mesophyll (in C<sub>3</sub> wheat) | Adding cyanobacterial HCO<sub>3</sub><sup>-</sup> transporters (single-subunit BicA and SbtA) to the chloroplast envelope (Price *et al.*, 2013). | A biochemical model of single-cell CCM photosynthesis is derived by combining the single-cell C<sub>4</sub> model (von Caemmerer, 2003) and the C<sub>4</sub> photosynthesis models (von Caemmerer, 2000) for modelling the cyanobacterium CCM in C<sub>3</sub>. Effects of adding cyanobacterial HCO<sub>3</sub><sup>-</sup> transporters are captured by:<br>• Additional cost of 0.75 ATP for active transport of HCO<sub>3</sub><sup>-</sup> into the mesophyll (Price *et al.*, 2011), which equates to 20% of the total ATP consumption (including the 3 ATP required by the C<sub>3</sub> cycle), to operate the two transporters; the HCO<sub>3</sub><sup>-</sup> transport rate with respect to CO<sub>2</sub> levels is aggregated in a single set of maximal activity ($V_{bmax}$) and Michaelis-Menten constant ($K_b = 60$); $V_{bmax}$ is assumed to scale linearly with leaf nitrogen content above the leaf structural N requirement for photosynthesis (slope = 0.36, which gives $V_{bmax}$ of 45 $\mu\text{mol m}^{-2} \text{s}^{-1}$ with wheat leaf SLN of 2 g N m<sup>-2</sup>;<br>• Induction of cyclic electron flow: in C<sub>4</sub> photosynthesis, in addition to the requirement by the C<sub>3</sub> cycle, the additional ATP costs in the mesophyll ($\phi$) induces cyclic electron flow. The same effect is modelled for CCM in C<sub>3</sub> by incorporating a multiplier $z$ ($= \frac{3-f_{cyc}}{4(1-f_{cyc})}$) to the linear electron flow ($J$), where $f_{cyc}$ is the fraction of $J$ out of PSI that proceeds via cyclic electron flow (von Caemmerer, 2021). This fraction is 0.25ϕ given that C<sub>4</sub> with the additional 2 ATP requirement has $f_{cyc}$ of 0.5 and C<sub>3</sub> with no additional ATP requirement has negligible $f_{cyc}$. The multiplier $z$ is calculated to be 0.75 when $\phi$ is negligible as in C<sub>3</sub> photosynthesis and is ca. 0.87 when $\phi = 0.75$ for operating the transporters.<br>• The original mesophyll conductance is modelled by the cell wall plasmalemma interface and chloroplast envelope conductance in series. They are assumed 1 and 1 mol m<sup>-2</sup> s<sup>-1</sup> bar<sup>-1</sup> (giving an intercellular–chloroplastic conductance of 0.5 mol m<sup>-2</sup> s<sup>-1</sup> bar<sup>-1</sup> |
| | | | with leaf SLN of $2 \text{ g N m}^{-2}$ at $25^\circ\text{C}$, comparable to that observed in $\text{C}_3$ wheat). |
| | (4.2) Cyanobacterial CCM: full implementation of the cyanobacterial CCM (in $\text{C}_3$ wheat) | Building on Target 4.1, the full CCM implementation involves adding carboxysomes containing carboxysomal Rubisco, and systems to further minimize $\text{CO}_2$ diffusion from the site of carboxylation, which are likely to involve eliminating carbonic anhydrase from the chloroplast stroma, adding NDH-1-based $\text{CO}_2$ pump, and/or making the chloroplast envelope less conductive to $\text{CO}_2$ via aquaporin manipulation (Price *et al.*, 2013). | The single-cell CCM photosynthesis model is capable of simulating the full CCM implementation. In addition to those changes described in Target 4.1, the full CCM effects are modelled by:<br>• Replacing endogenous wheat Rubisco with a carboxysomal Rubisco. Kinetic properties of a carboxysomes-encapsulated Rubisco with $K_c$ increased by a factor of 9.9; $K_o$ reduced by 17%, and $S_{c/o}$ is reduced by 39%; derived from comparing Rubisco kinetics reported for wheat (Sharwood *et al.*, 2016b) and *Cyanobium* Rubisco produced in transgenic tobacco plants (Long *et al.*, 2018); assuming carboxylation and oxygenation temperature responses remain similar.<br>• Chloroplast envelope conductance is expected to reduce to levels similar to that in $\text{C}_4$ species (i.e. $3 \text{ mmol m}^{-2} \text{ s}^{-1}$) (von Caemmerer, 2000)) with the installation of carboxysomes (having the desirable properties of retarding $\text{CO}_2$ leakage and negligible effects on entry and exit of $\text{HCO}_3^-$) together with the additional biochemical changes regarding carbonic anhydrase and NDH-1-based $\text{CO}_2$ pump.<br>• Assuming no change in $V_{\text{cmax}}$ and photosynthetic machinery N requirement with Rubisco replacement and carboxysome synthesis. $k_{\text{cat}}^c$ of the encapsulated Rubisco could theoretically attain the levels of free *Cyanobium* Rubisco produced in the tobacco plants and those from *Cyanobium* cells (ca. $9.5 \text{ s}^{-1}$), which is a factor of up to ca. 3 compared to those of wheat (Long *et al.*, 2018). This means the same $V_{\text{cmax}}$ can be achieved with less Rubisco, resulting in no additional N requirement or surplus N even after the costs of carboxysome are taken into account (Rae *et al.*, 2017).<br>• Assuming efficient bicarbonate transport systems to achieve sufficiently high levels of chloroplastic bicarbonate concentrations. This is modelled by setting the maximal activity of the $\text{HCO}_3^-$ transporter ($V_{\text{bmax}}$) at a level comparable to the maximal PEP carboxylase activity used in $\text{C}_4$ sorghum (i.e. $V_{\text{bmax}}$ of $\sim 120 \mu\text{mol CO}_2 \text{ m}^{-2} \text{ s}^{-1}$ with wheat leaf SLN of $2 \text{ g N m}^{-2}$ at $25^\circ\text{C}$); if achieved by having more of both BicA and SbtA there would be no change in $K_b$; temperature response of $V_{\text{bmax}}$ and $K_b$ are assumed same as the $\text{C}_4$ equivalent in the initial carboxylation step. |
| | (5) Higher phosphoenolpyruvate carboxylation rate (in C<sub>4</sub> sorghum) | Over expressing PEP carboxylase (von Caemmerer & Furbank, 2016) | +20% maximum carboxylation rate of phosphoenolpyruvate ($V_{pmax}$) |
|  | (6) Reduced CO<sub>2</sub> leakage from bundle sheath (in C<sub>4</sub> sorghum) | Manipulating the interface between mesophyll and bundle sheath | –20% bundle sheath conductance for gases, affecting efflux of CO<sub>2</sub> and O<sub>2</sub> from the bundle sheath to the mesophyll, without affecting diffusive flux of the C<sub>4</sub> and C<sub>3</sub> cycle photosynthetic metabolites between the cells. |
| Electron transport chain | (7) More efficient electron transport chain | Overexpressing the Rieske-FeS protein to generate more abundance of cytochrome b<sub>6</sub>f complex (Ermakova *et al.*, 2019, Simkin *et al.*, 2017). | Average of effects on parameters describing the $J-I$ response inferred from photosynthetic light response data by Ermakova *et al.* (2019) and Simkin *et al.* (2017): +20% $J_{max}$ (maximum electron flow at saturating light); +15% $J$ response at low light; –40% empirical $J-I$ curvature factor; assuming $J_{max}$ has the same temperature response. |
|  | (8) Access to additional light energy from photons in the 700–750 nm wavelength; increasing the fraction of PAR to total solar. | Swapping some chlorophyll *a* and *b* in Photosystems II and I with cyanobacterial Chlorophyll *d* and *f* by in all leaves of the canopy (Chen & Blankenship, 2011). | +20% photosynthetic active radiation from solar radiation (both direct and diffuse) to all leaves in the canopy assuming Chlorophyll *d* and *f* can be constitutively expressed in all leaves and they are able to capture all photons in the 700–750 nm wavelength (Chen & Blankenship, 2011). Assumed no change to plant's ability to use the visible spectrum (400–700 nm); electron flow has the same light and temperature responses. |
| Manipulation stacking | (9) Combining “better” Rubisco, higher electron transport rate, and improved diffusion of CO<sub>2</sub> into the mesophyll | Combinations of the bioengineering approaches described above. | Combining parameter changes described in Targets 2, 3 and 7. |

### Leaf and canopy photosynthesis simulation

Simulation of *A*–*C*_i_ was performed using the diurnal canopy photosynthesis modelling framework (Wu *et al*., 2018, Wu *et al*., 2019) expanded to include a generic photosynthesis-CO_2_ diffusion model that can be switched between the C_3_, C_4_ and the (single-cell design) cyanobacterial CCM pathways depending on settings for calculated variables (Appendix A: Table S2). The baseline set of the C_3_ and CCM wheat, and C_4_ sorghum photosynthesis model parameters were adapted from Wu *et al*. (2019) with some recalculated using new data (Table S5). Key physiological parameters are the maximum rate of Rubisco carboxylation (*V*_cmax_), maximum rate of PEP carboxylation (*V*_pmax_), and maximum rate of electron transport at infinite light intensity (*J*_max_). The baseline values of *V*_cmax25_ and *J*_max25_ for wheat (the subscripted number denotes value at the standard 25°C), and *V*_cmax25_, *V*_pmax25_ and *J*_max25_ for sorghum were set to those observed previously (Silva-Pérez *et al*., 2017, Sonawane & Cousins, 2020, Sonawane *et al*., 2017). The value of *J*_max25_ along with *α*_PSII_ and *θ* used in Eqn S4 gave a potential whole-chain linear electron transport rate (*J*) of 232 μmol m^−2^ s^−1^ at photosynthetic photon flux density (PPFD) of 1800 μmol m^−2^ s^−1^ and 25°C, comparable to that inferred from C_3_ wheat data (Silva-Pérez *et al*., 2017). The ATP-limited version of the electron transport-limited equation was used in the single-cell CCM model with a factor that relates *J* to the production of ATP (Eqns S11, S13) following von Caemmerer (2021). This treatment gave almost the same electron-transport-limited CO_2_ assimilation rate to the NADPH-limited equation used in the C_3_ model (Eqn S2). The C_4_ model also uses the ATP-limited version of the equation. For the C_4_ electron transport parameters, the value of *J*_max25_ along with *α*_PSII_ and *θ* gave a *J* of 215 μmol m^−2^ s^−1^ at PPFD of 1800 μmol m^−2^ s^−1^ and 25°C comparable to that inferred from C_4_ maize data (Massad *et al*., 2007). The maximal activity of the bicarbonate transporters (*V*_bmax_) was taken from Price *et al*. (2011). In the full CCM case, a more efficient CO_2_ transportation rate comparable to that in the C_4_ version of the CCM was used as the system would require a higher inorganic carbon influx to function efficiently. If a low *V*_bmax_ was used, yield would be significantly impacted due to reduced CO_2_ assimilation rate and growth (Figure S9).

The key physiological parameter (i.e. *V*_cmax25_, *J*_max25_, and *V*_pmax25_) values were used to calculate the corresponding *χ* values for input into the cross-scale model, where each *χ* value is the slope of the linear relationship between the photosynthetic parameter and specific leaf nitrogen (SLN, g N m^−2^ leaf) (Table S5). The Rubisco catalytic properties and mesophyll conductance, the C_4_ bundle sheath conductance, and the baseline *C*_i_/*C*_a_ were taken from published data (Bernacchi *et al*., 2002, Boyd *et al*., 2015, Long *et al*., 2018, Massad *et al*., 2007, Ubierna *et al*., 2017, von Caemmerer & Evans, 2015) and a summary table by Wu *et al*. (2019). The Michaelis-Menten constant for CO_2_ (converted from the constant for bicarbonate) (*K*_b_) (Price *et al*., 2011) was calculated from the CO_2_ response resulting from BicA and SbtA transporters combined. The *V*_bmax_ value was the sum of the two transporters using values from Price *et al*. (2011). The photosynthetic parameters in Table S5 were used for simulating the baseline C_3_ and C_4_ *A*_c_ and *A*_j_ limitations, and *A*–*C*_i_ curves (Figures 2 and 3). The curves were comparable to those observed previously (Silva-Pérez *et al*., 2017, Sonawane *et al*., 2017).

**Figure 1.**
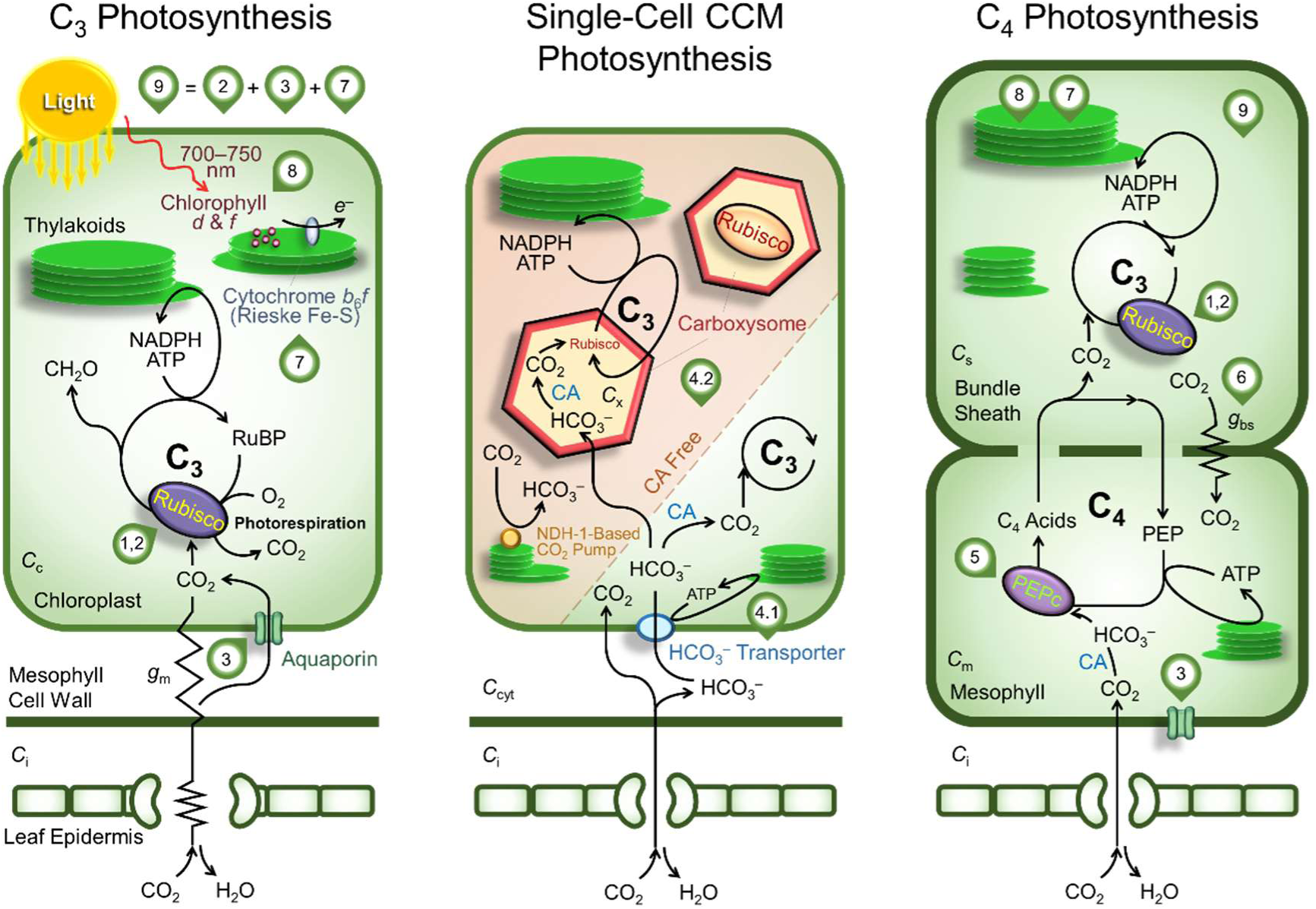
Overview of the leaf photosynthetic pathways and manipulation targets used in wheat and sorghum crop growth and yield simulations. The manipulation targets are numbered here and detailed in Table 1. Bioengineering strategy “1,2” encompasses “1.1”, “1.2”, “1.3”, “1.4”, and “2”. Strategy “9” is achieved by stacking “2”, “3”, and “7”. Graphics for stomatal and mesophyll resistance/conductance are omitted in the single-cell CCM and C_4_ photosynthesis pathways for simplicity. Abbreviations: *C*_i_, *C*_c_, *C*_m_, C_x_, and *C*_s_ are the intercellular, chloroplastic, mesophyll, carboxysomal, and bundle sheath CO_2_ partial pressures; CA, carbonic anhydrase; *g*_bs_, bundle sheath conductance; *g*_m_, mesophyll conductance.

**Figure 2.**
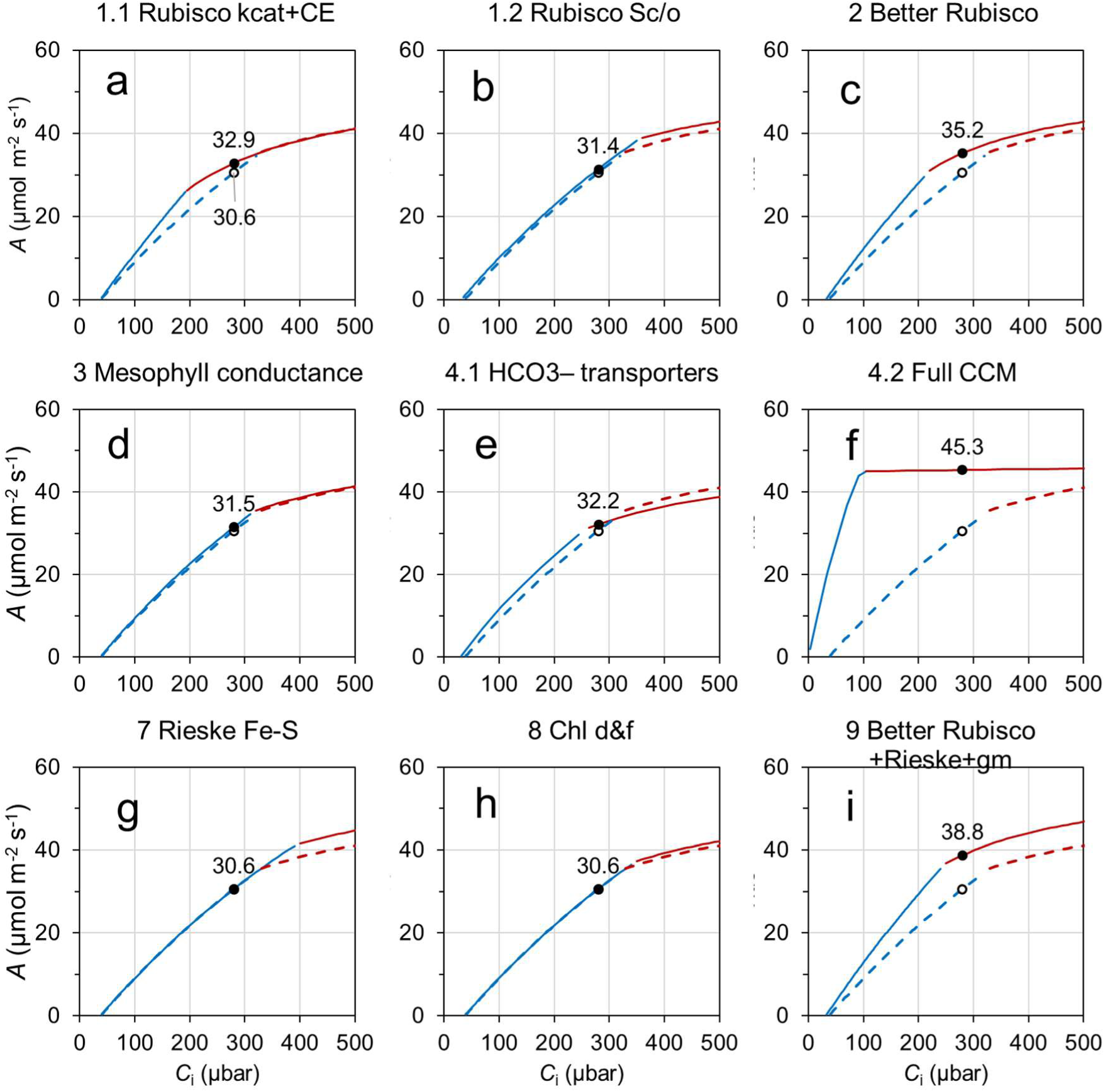
Simulated C_3_ wheat leaf photosynthetic response to intercellular CO_2_ (*A–C*_i_) for the baseline and manipulated scenarios. *A–C*_i_ are simulated for 25°C with photosynthetic photon flux density of 1800 μmol m^−2^ s^−1^ using the C_3_ and single-cell CCM photosynthesis model parameter values given in Table S5. Panels are for the different leaf photosynthetic manipulations as described in Table 1. The baseline *A–C*_i_ is reproduced in every panel as dashed lines; solid lines are *A–C*_i_ with photosynthetic manipulation. Blue and red are Rubisco activity (*A*_c_) and electron transport (*A*_j_) limited *A*, respectively. Un-filled and filled circles are *A* at an ambient CO_2_ of 400 μbar (i.e. intercellular CO_2_ of 280 μbar) for the baseline and with manipulations, respectively. The value of the baseline *A* is indicated in Panel (a); the manipulated *A* is given in all panels. (a–c) relate to Rubisco function manipulations, (d–f) relate to CO_2_ delivery manipulations, and (g–h) relate to electron transport chain manipulations, (i) a combination of the three aspects. Details of the manipulations are given in Table 1.

**Figure 3.**
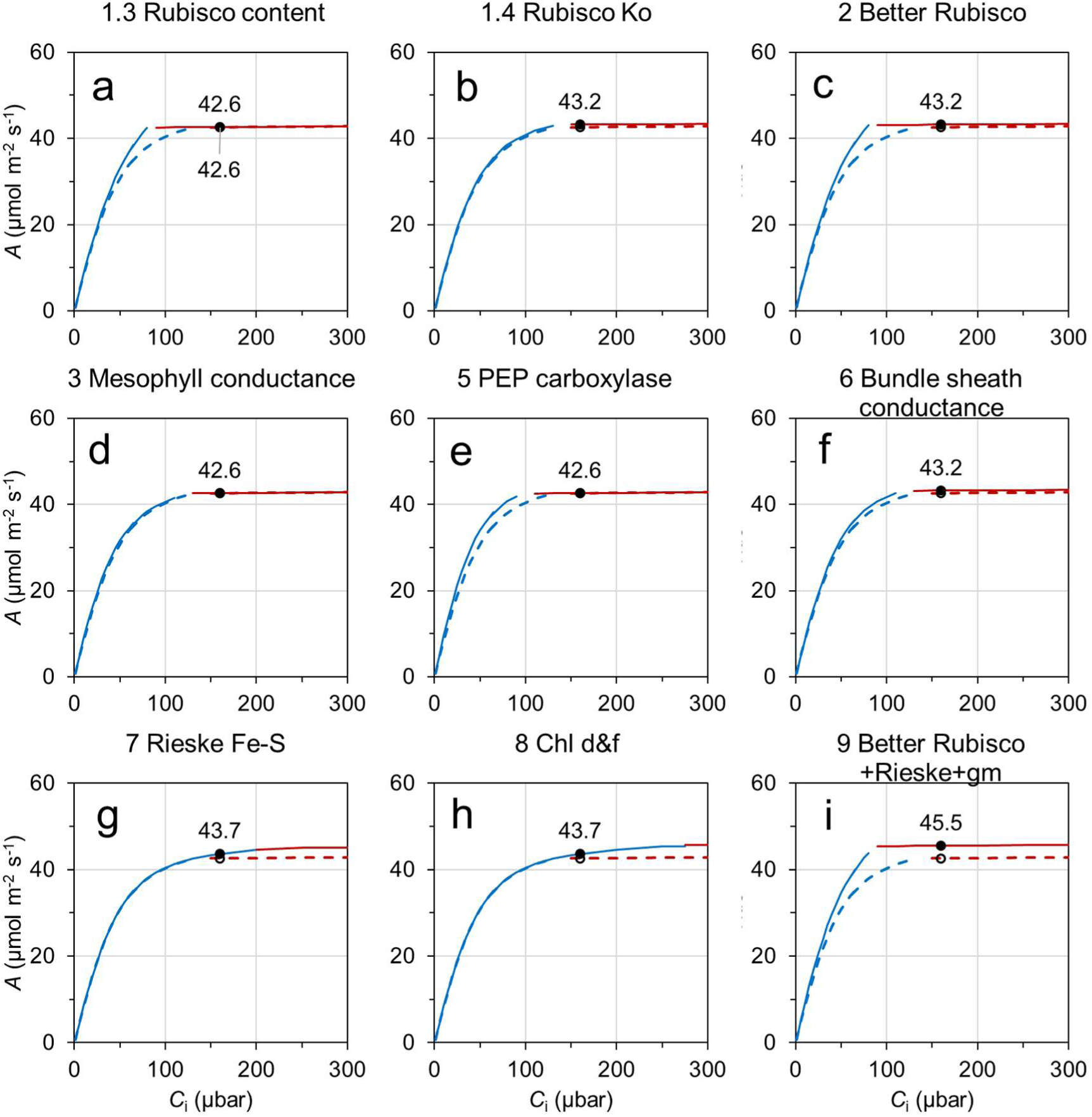
Simulated C_4_ sorghum leaf *A–C*_i_ for the baseline and manipulated scenarios. *A–C*_i_ is simulated for 30°C with photosynthetic photon flux density of 1800 μmol m^−2^ s^−1^ using the C_4_ sorghum photosynthesis model parameter values given in Table S5. Lines and symbols are the same as those described in Figure 2.

The diurnal canopy photosynthesis modelling approach used here calculates canopy photosynthesis by partitioning canopy leaf area into sunlit and shaded fractions (on a per ground area basis), integrates the key photosynthetic parameters over the respective leaf fraction over the ground area, and calculates the *A*_c_ and *A*_j_ on a fraction basis (Wu *et al*., 2018). Unlike the leaf-level *A*_c_ and *A*_j_, the fraction-level *A*_c_ and *A*_j_ represent the collective rates of all leaves in the fraction, having incorporated within canopy variations in intercepted light and photosynthetic parameters through canopy depth. The model assumes photosynthesis and stomatal conductance respond instantaneously to changing light conditions and attain steady-state levels (Wu *et al*., 2018). This approach was demonstrated to predict field-observed crop growth rates well over the period of a crop cycle (Wu *et al*., 2019). Fraction-level *A*_c_ and *A*_j_ can also be plotted to give *A*–*C*_i_ curves (e.g. Figures S1 and S2).

### Modelling leaf photosynthetic manipulation

A broad list of manipulation strategies is examined in this study. A comprehensive description on how each strategy is theorised to function is given in Table 1 with full references to findings from transgenic and previous modelling studies. The manipulations include enhancing Rubisco function by enhancing its catalytic properties and/or content (manipulation outcomes 1.1, 1.2, 1.3, 1.4) (Martin-Avila *et al*., 2020, Salesse-Smith *et al*., 2018, Sharwood *et al*., 2016a, Sharwood *et al*., 2016b); a “better” Rubisco from stacking Rubisco function enhancements (outcome 2); enhancing CO_2_ delivery by improving mesophyll conductance (outcome 3) (Groszmann *et al*., 2017), installation of a cyanobacterial CO_2_ concentrating mechanism in C_3_ wheat (outcomes 4.1 and 4.2) (Price *et al*., 2013), overexpression of PEP carboxylase in C_4_ sorghum (outcome 5), or reducing bundle sheath conductance in C_4_ sorghum (outcome 6); enhancing electron transport rate by overexpression of the Rieske-FeS protein of the cytochrome *b*_6_f complex (outcome 7) (Ermakova *et al*., 2019, Simkin *et al*., 2017), or extending useful photosynthetically active radiation to 700–750 nm of leaves by supplementing light-harvesting complexes with cyanobacterial Chlorophyll *d* and *f* in all leaves of the canopy (outcome 8) (Chen & Blankenship, 2011). A tangible case of stacking a selection of some of these strategies was also included (outcome 9: “better” Rubisco, overexpression of Rieske-FeS protein, and improved mesophyll conductance).

Nitrogen costs of achieving manipulation outcomes can be assumed neutral. Modifying Rubisco kinetic properties (outcomes 1.1, 1.2, 1.4) and swapping chlorophyll types (outcome 8) have minimal net N cost requirement. N cost associated with increased expression of proteins for manipulation outcomes 3, 4.1, 5, 6, and 7 is likely to be small (Evans & Clarke, 2019). Increasing Rubisco content in C_4_ sorghum (outcomes 1.3 and 2) is also likely small in N cost due to a lower baseline content. Additional N cost associated with both bicarbonate transporters and whole carboxysomes (outcome 4.2) could be offset by savings from reduction in Rubisco content as detailed in Table 1 (Rae *et al*., 2017). Therefore, it was assumed that photosynthetic manipulations were achieved with no effects in N demand of expanding leaf, leaf structural N requirement (or minimum leaf N), and N translocation from leaves to other plant organs (van Oosterom *et al*., 2010a,b).

The C_3_ photosynthesis setting of the photosynthesis-CO_2_ diffusion model was used for most of the wheat photosynthetic manipulation simulations, except that the single-cell CCM setting was used to model installation of the CCM. The C_4_ setting was used for all of the sorghum photosynthetic manipulations. Manipulations have different effects on the *A*_c_ and *<u>A</u>*_j_. Examples of predicted consequences of these manipulations on and limitations and leaf-level *A*–*C*_i_ curves are shown in Figures 2 and 3.

### Dynamic crop growth and yield simulation

Multiyear × location crop growth simulations, akin to extensive multi-environment trials, were conducted using common wheat and sorghum cultivars to understand and quantify consequences of leaf photosynthetic manipulation on crop growth and yield over a wide range of environments. This involved running simulations with representative daily weather data at selected sites across crop production regions. Australian environments were used in this study as the year-to-year environmental condition variability present a diverse set of non-stressed and stressed conditions and can generate a wide range of yield levels. The median sowing date, median amount of stored soil water at sowing, and the most commonly used agronomy and N application for the crop were used in this multi-environment simulation (Table S4).

The weather and soil aspects of the simulations were parameterised depending on the crop in question and the production site. The target population of environments for wheat in Australia has been classified into six distinct types based on a principal component analysis of long-term year-to-year production variability at shire scale (Potgieter *et al*., 2002) (Figure S3). One production site representative of each of these six regions was selected based on its loading for the respective principal component as well as being a key centre/town for wheat production (Table S4). Similar considerations were followed in selecting the four sites from north-eastern Australia for sorghum production simulation.

Interannual weather variability at each site was represented by accessing its long-term (1900-2020) daily weather record (including maximum and minimum air temperature, incoming solar radiation, and precipitation), which was obtained from the SILO patched point data set (http://www.longpaddock.qld.gov.au/silo/index.html; Jeffrey *et al*. (2001)). The intention was not to simulate historical yield levels, but to use historical weather data to sample interannual weather variabilities. Ambient CO_2_ was set at 400 ppm (*ca*. 400 μbar). Detailed parameterisations of soil characteristics (including soil depth, plant available water capacity and typical N present in the soil at sowing) were taken from Chenu *et al*. (2013) and Hammer *et al*. (2014).

Medium-maturing wheat (Janz) and sorghum (Hybird MR-Buster) cultivars were used in the multiyear × location simulation (Table S4). Their physiology reflects the commonly used cultivars in Australian production environments and their physiological response to environmental variables have been well-parameterised in APSIM crop growth models and tested (Ababaei & Chenu, 2020, Hammer *et al*., 2010).

Locally adapted agronomic practices for the different sites were used. Briefly, wheat is sown around May–June each year, while sorghum has a wider sowing window between October and January. Sowing dates used in this multiyear × location simulation were the median values calculated from the reported uniform distribution of dates within the sowing windows (Ababaei & Chenu, 2020, Hammer *et al*., 2014). A row-planting configuration was used for both crops with sorghum having a 1–meter row spacing and 5 plants m^−2^, while wheat had 0.25–meter row spacing and a density of 100 or 150 plants m^−2^ (Table S4). Starting soil water content was set to the median values, which were calculated from the frequencies reported for wheat (Chenu *et al*., 2013) and sorghum (Hammer *et al*., 2014). Soil N at time of sowing ranged between 30 and 50 kg ha^−1^. The sorghum crop is typically fertilized with N before or at sowing with N applied to the surface soil layers, while for wheat N application can also occur later in the growing season depending on crop stage and soil water/precipitation conditions (Table S4). The weather variability, crop configuration, and N application combinations present a broad spectrum of non-stressed to stressed production conditions.

During each crop growth simulation cycle the cross-scale model simulated interactions between growth/photosynthesis, light interception and water use, crop development, resource (water, nitrogen) supply–demand balance, carbohydrate and nitrogen allocation among organs, growth of grains, and effects of environmental variables (sunlight, water, temperature, and nitrogen). Each crop growth simulation involved hourly canopy photosynthesis simulation over the diurnal period from early in the crop cycle (with leaf area index ≥ 0.5) to physiological maturity. Trajectories of simulated crop attributes through the crop cycle were extracted for detailed analysis (e.g. Figures 4 and S4). The plots exemplify a medium-yielding wheat and sorghum crop from the set of 120 seasons of the baseline simulation at the Dalby site with the median sowing date and starting soil water.

**Figure 4.**
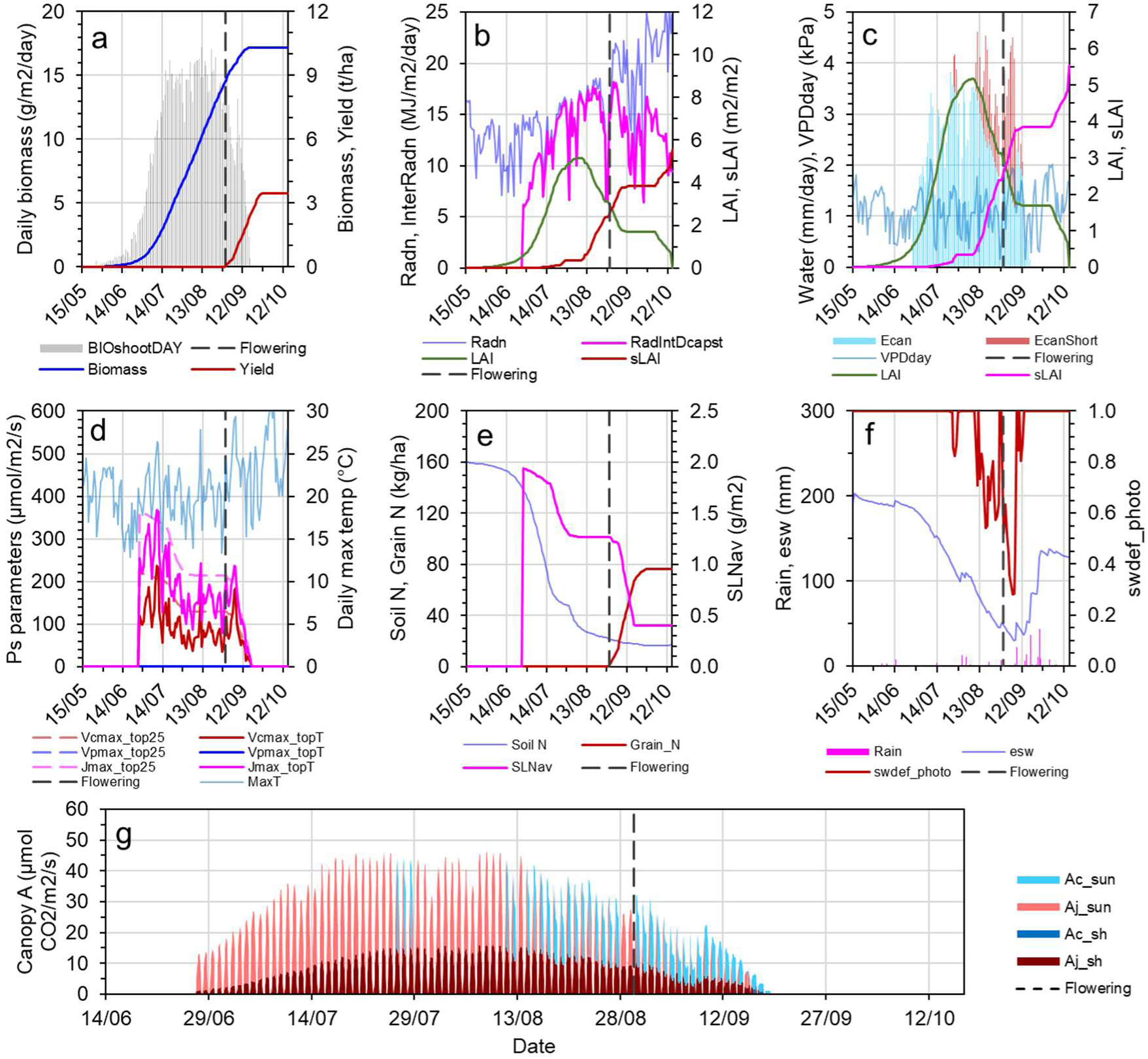
Predicted wheat crop attributes dynamics, and environmental variables over a sample crop cycle. Results are from a medium-yielding year at the Dalby site with the medium sowing date and starting soil water (Table S4). (a) Cumulative crop biomass and yield. (b) Canopy leaf area index, solar radiation and interception. (c) Potential crop water demand is shown by the bars, which is made up of a fraction that is met by supply from soil water uptake by roots (i.e. actual water use) and a fraction that is not met (red bars). (d) Photosynthetic parameters for the uppermost leaves of the canopy at 25°C and the maximum air temperature during the day. (e) Soil N supply and crop N status including specific leaf nitrogen and N in grains. (f) Plant extractable soil water and a crop water stress factor; a value of 1 means all crop water demand is being met, while 0 means no water is available. (g) Daily canopy photosynthesis; each peak is made up of a histogram of total canopy photosynthesis on an hourly timestep over one diurnal period. An equivalent figure for sorghum is shown in Figure S4. Abbreviations: BIO_shootDAY_, daily shoot biomass growth; Radn, daily incident solar radiation, RadIntDcapst, daily intercepted radiation by the whole canopy; LAI, leaf area index; sLAI, senescenced LAI; Ecan, actual crop water use; EcanShort, fraction of the potential demand not met by supply; VPDday, indicative daytime vapour pressure deficit; Vcmax_top25, Vpmax_top25, Jmax_top25 are the values of the maximum rate of Rubisco carboxylation, maximum rate of PEP carboxylation, and maximum rate of electron transport at infinite light at 25°C; Vcmax_topT, Vpmax_topT, Jmax_topT are those photosynthetic parameter values calculated using the maximum temperature of the day (MaxT); SLNav, canopy-average specific leaf nitrogen; esw, plant extractable soil water; swdef_photo, a crop water stress factor given by EcanFilled divided by the sum of EcanFilled and ECanShort; Ac_sun and Aj_sun, Rubisco activity and electron transport limited gross CO_2_ assimilation rate of the sunlit fraction of the canopy (only the lower of the two limitations is shown); Ac_sh and Aj_sh, the same limitations for the shaded fraction.

Changes in dynamics of the crop attributes and final grain yield were also predicted for the different photosynthetic manipulation strategies across the locations using the same sowing date and starting soil water (Table S4). Crop attribute trajectories with and without photosynthetic manipulation were generated and used in detailed analysis (e.g. Figures S5– S7). Consequences of photosynthetic manipulations for grain yield were quantified using change in simulated yield relative to the baseline parameterisation across the range of production environments in this multiyear × location simulation (Figures 5 and 6). The yield change associated with photosynthetic manipulation for each simulation crop-year was plotted against yield level for the baseline scenario. Quantile regression was performed in Python using the statsmodels’ QuantReg class to identify the 10^th^ and 90^th^ percentile regressions in the plots to delineate the upper and lower percentage yield change (Figures 5 and 6).

**Figure 5.**
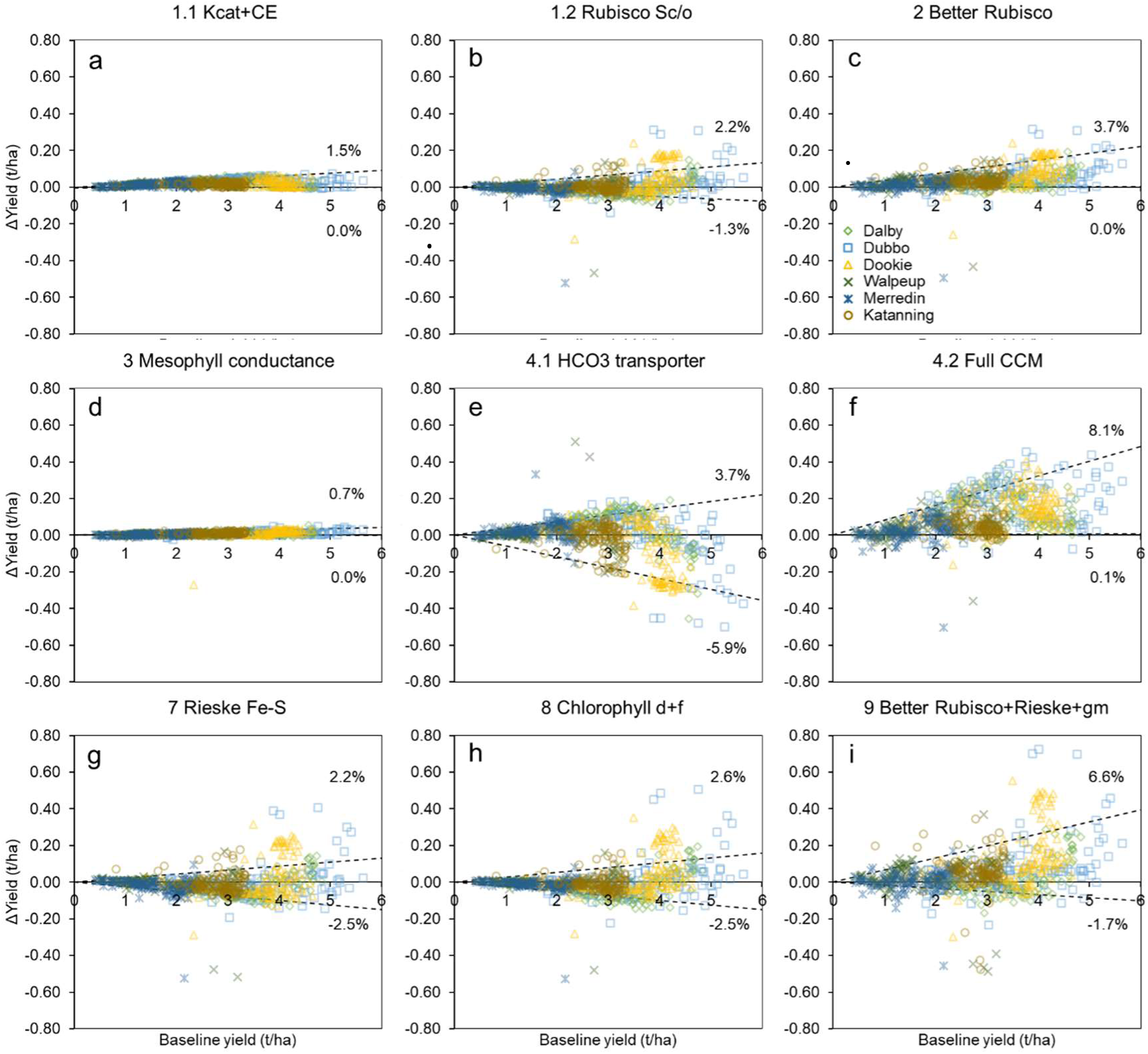
Predicted change in wheat yield (t/ha) relative to the baseline simulations for leaf photosynthetic manipulations. Panels give results for the different manipulation strategies (Table 1); results are plotted together for the six contrasting sites across the Australian wheatbelt. This focused set of simulation uses representative seasonal weather data sampled from the past 120 years (1900-2020), the medium sowing date, and plant available water at sowing specific for each site (Table S4). The dashed lines indicate the 10^th^ and 90^th^ percentile regressions for Δyield versus baseline yield. Their slopes indicate the upper and lower percentage yield changes (n = 1,440 crop cycles per panel; 720 baseline and 720 with manipulation).

**Figure 6.**
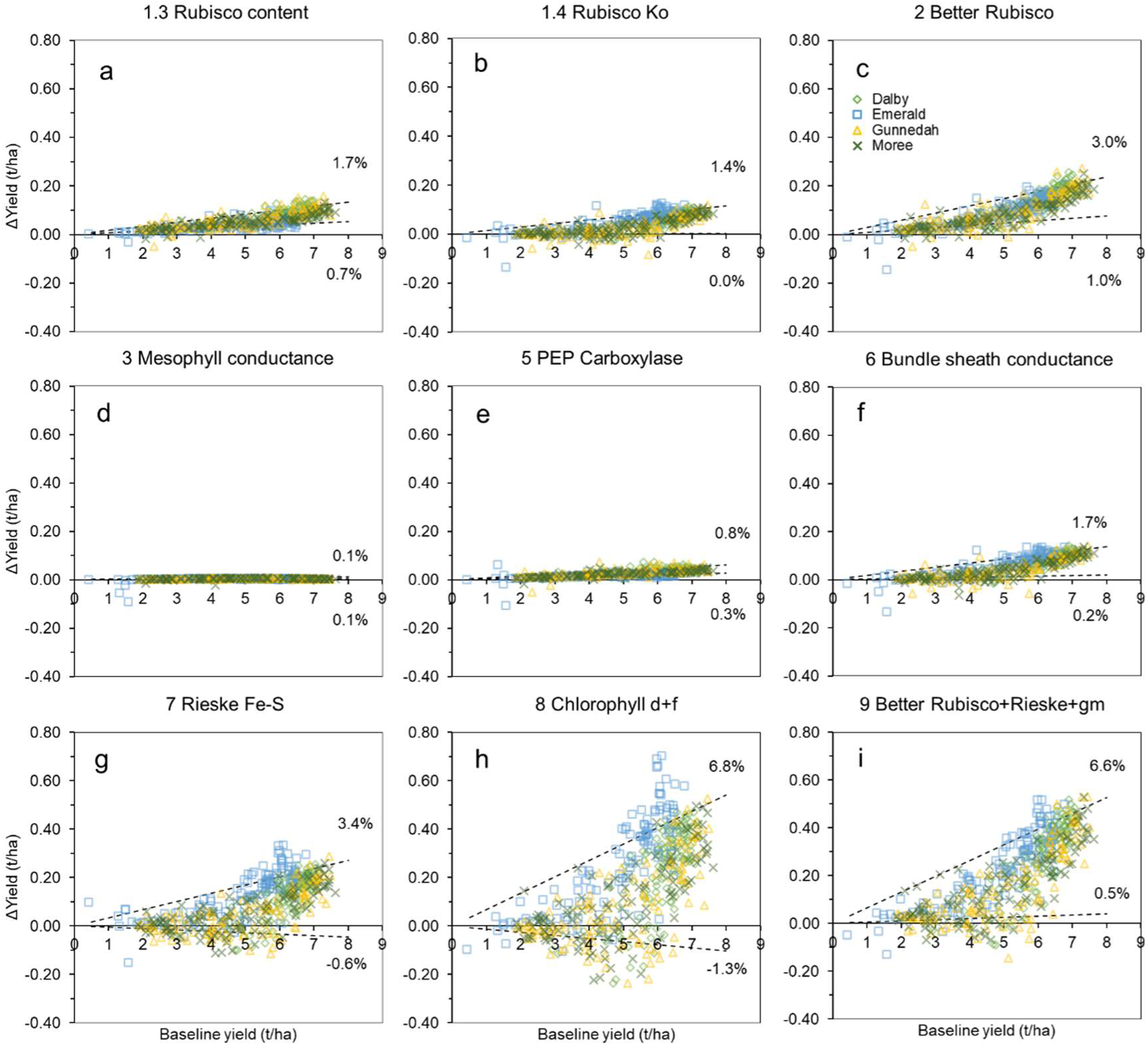
Same as for Figure 5 for predicted sorghum yield changes. Results are plotted together for the four contrasting sites across sorghum production regions (n = 960 crop cycles per panel).

### Australian crop production simulation

As a case study, the consequences for national scale crop production of photosynthetic manipulations were quantified by using baseline production at the regional scale combined with the extent of impact of each leaf photosynthetic manipulation strategy. Historical Australian wheat (1901-2004) and sorghum (1983-2015) production data at the regional level (Potgieter *et al*., 2002) were averaged and used as the baseline (Table S6). For quantifying yield impact of the manipulations, the multiyear × location simulation was expanded to including sowing dates and starting soil water levels as additional factors. Three representative levels of each of sowing date and starting soil water were calculated from their distributions (as described above) and used the in the simulation (Table S4). The overall year × sowing date × soil water × site × manipulation amounted to 194k crop cycles, for wheat and sorghum combined. This ensured balanced representation of all possible starting conditions at sowing in the crop production simulations. The median percentage change in grain yield, and the first and third quartile values at each representative site (Table S7) were predicted and applied to the corresponding regional scale production. National scale impact was calculated by weighting regional contribution to national production (Table S6).

## Results and Discussion

While understanding and bioengineering of photosynthetic pathways have advanced significantly over the past decades with promising evidence at the leaf level, however, assessment at the crop level across contrasting production environments remains limited or indirect. This hampers application of photosynthesis research in crop improvement (Fischer *et al*., 2014). In this study, we used an advanced cross-scale crop growth model (Wu *et al*., 2019) to generate novel understanding in how potential photosynthetic manipulation can affect crop growth and yield dynamics in a wide range of environments. Predictions on leaf and canopy, to crop growth and yield are presented and discussed.

### Changes in leaf and canopy photosynthesis with manipulations

The leaf-level photosynthetic response to intercellular CO_2_ (*A–C*_i_) with and without (baseline scenario) manipulations were predicted for C_3_ wheat and C_4_ sorghum using the photosynthetic parameter values in Table S5 and compared with prior knowledge. For C_3_ wheat at a photosynthetic photon flux density (PPFD) of 1800 μmol m^−2^ s^−1^ and 25°C, *A* was enzyme limited (*A*_c_) at low *C*_i_ and electron transport limited (*A*_j_) at high *C*_i_ (Figure 2). Transition from *A*_c_ to *A*_j_ occurred slightly above *C*_i_ = 300 μbar suggesting *A*_c_ limitation at ambient CO_2_ (i.e. *C*_i_ = 280 μbar using *C*_a_ = 400 μbar and *C*_i_/*C*_a_ = 0.7). For C_4_ sorghum at a PPFD of 1800 μmol m^−2^ s^−1^ and 30°C, *A*–*C*_i_ showed a steep *A*_c_-limited initial CO_2_ response below a *C*_i_ of ∼125 μbar followed by *A*_j_ limitation above that *C*_i_ (Figure 3). Thus *A* was limited by *A*_j_ at ambient CO_2_ (i.e. *C*_i_ = 160 μbar with *C*_i_/*C*_a_ of 0.4). This is consistent with evidence that electron transport can limit C_4_ photosynthesis under high-light conditions (Ermakova *et al*., 2019). The simulated baseline *A*–*C*_i_ for wheat and sorghum were comparable to previously published data (Silva-Pérez *et al*., 2017, Sonawane *et al*., 2017).

Rubisco function manipulations were predicted to predominantly affect *A*_c_ at low *C*_i_ (Figures 2a–c, 3a–c). The CO_2_ delivery related manipulations affected both *A*_c_ and *A*_j_ (Figures 2d–f, 3d–f). The electron transport chain related manipulations affected *A*_j_ at high *C*_i_ (Figures 2g–h, 3g–h). Stacking all three aspects affected both *A*_c_ and *A*_j_ (Figures 2i, 3i). Specifically, manipulation of C_3_ wheat Rubisco carboxylation rate and carboxylation efficiency to achieve those of C_4_ maize values (Table 1: outcome 1.1) was predicted to improve *A*_c_ and lower the *C*_i_ of the *A*_c_–*A*_j_ limitation transition (Figure 2a), which are consistent with previous simulation analysis (Sharwood *et al*., 2016b). Enhancement of wheat Rubisco specificity for CO_2_ improved both *A*_c_ and *A*_j_, but more so for the latter (Figure 2b). Manipulation of C_4_ Rubisco improved *A*_c_ (Figure 3a), comparable to observations in maize transgenics with increased Rubisco content (Salesse-Smith *et al*., 2018). The combination of the enhancement of Rubisco properties (i.e. the “better” Rubisco) (Table 1: outcome 2) generated an additive effect of the component enhancements in both wheat and sorghum (Figures 2c, 3c).

On the CO_2_ delivery related manipulations, increasing mesophyll conductance (Table 1: outcome 3) had minimal impact on *A*–*C*_i_ response in both wheat and sorghum (Figures 2d, 3d). A previous simulation of photosynthetic CO2 assimilation rate across a wide range of mesophyll conductance values had also shown little effect on *A* unless mesophyll conductance was low (Groszmann *et al*., 2017). Installing the cyanobacterial bicarbonate transporters alone (Table 1: outcome 4.1) was predicted to improve *A*_c_ (Figure 2e) as the active transport mechanism elevated CO_2_ level at the site of Rubisco carboxylation. This was consistent with previous modelling results of *HCO*_1_ transporter addition to C_3_ photosynthesis (Price *et al*., 2011). However, a reduction in *A*_j_ was predicted as the elevated CO_2_ could not compensate for the extra ATP requirement of the bicarbonate transporters (Figure 2e). The installation of the full cyanobacterial CCM (Table 1: outcome 4.2) was predicted to generate the greatest changes in the *A*–*C*_i_ response (Figure 2f). This extent of effect agreed with a previously study using a more elaborate model of a CCM (McGrath & Long, 2014). In C_4_ sorghum, the CO_2_ delivery related manipulations (Table 1: outcomes 5 and 6) affected *A*_c_ with smaller changes in *A*_j_ (Figure 3d,e,f).

Predicted C_3_ and C_4_ *A*–*C*_i_ with Rieske-FeS protein overexpression (Table 1: outcome 7) increased in *A*_j_ and reflected responses observed experimentally in transgenic plants (Ermakova *et al*., 2019, Simkin *et al*., 2017) (Figures 2g, 3g). Addition of chlorophyll *d* and *f* (Table 1: outcome 8) had only limited effect on *A*_j_ in C_3_ wheat as electron transport rate was near saturation under the high-light condition used (Figure 2h). A larger effect on *A*_j_ was predicted for C_4_ sorghum (Figure 3h), which is consistent with a higher light saturation point in C_4_ photosynthesis (Ermakova *et al*., 2019). The combination of the “better” Rubisco, Rieske Fe-S protein, and mesophyll conductance (Table 1: outcome 9) was predicted to increase both *A*_c_ and *A*_j_ in both wheat and sorghum (Figures 2i, 3i).

It is important to note that demonstrating increases in CO_2_ assimilation rates using some specific conditions is not sufficient for understanding crop growth and yield consequences. Effect of manipulation strategies on CO_2_ assimilation rate needs to be assessed against many factors. These include changes in the incident solar radiation due to the relative movement of the sun across the sky and air temperature across the growing season. In addition, it is the photosynthesis of the whole canopy that drives crop biomass growth, which is influenced by canopy leaf area index (LAI, m^−2^ leaf m^−2^ ground) and specific leaf N (SLN, g N m^-2^ leaf), both of which change throughout the crop life cycle. The diurnal canopy photosynthesis modelling approach used here calculates canopy photosynthesis by predicting and summing CO_2_ assimilation rates of the sunlit and shaded leaf area fractions of the canopy as described in the Methods. Exemplary sunlit-fraction *A*–*C*_i_ are shown in Figures S1 and S2. Relative to the leaf level, the sunlit-fraction *A*–*C*_i_ has higher *A*_c_ and *A*_j_ due to integration of the enzyme-limited and electron transport-limited rates over its leaf area. However, *A*_c_ typically increases more relative to *A*_j_ and causes a reduction in the transition *C*_i_ (compare Figures S1 and S2 with Figures 2 and 3). This occurs because incident PPFD on a leaf area basis does not scale linearly with the leaf area of the sunlit fraction due to leaf orientations in a crop canopy. Therefore, the *A*_c_–*A*_j_ transition for the sunlit fraction would shift to lower *C*_i_ (e.g. compare Figures 2a). The shaded-fraction *A*–*C*_i_ would be dominated by *A*_j_ limitation due to low incident PPFD.

Photosynthetic manipulation effects on fraction-level *A*_c_ and *A*_j_ were comparable, in relative terms, to those described for the leaf level (Figures S1, S2). The interactions between the *A*_c_, *A*_j_, operating *C*_i_, environmental conditions, and canopy status, underpin the dynamics of canopy photosynthesis, stomatal conductance/crop water use, and these determine crop growth and resource demands over the crop cycle as discussed below. The effect of water stress is simulated by restricting stomatal conductance calculated by the Penman-Monteith Combination equation (Wu *et al*., 2019), in which case the operating *C*_i_ would be reduced, thus leading to reduced *A* and possible limitation by *A*_c_ (e.g. Figure S1a). Under limited transpiration and reduced stomatal conductance, *A*_c_ enhancement can still increase *A* by reducing *C*_i_ and improves intrinsic water use efficiency (e.g. Figure S1a). This suggests benefit of *A*_c_ enhancement is larger when water limitation is affecting photosynthesis. *A*_j_ enhancement is more relevant and beneficial without water limitation and when stomatal conductance can increase with enhanced CO_2_ assimilation rate (e.g. Figure S1i). However, the higher stomatal conductance drives higher transpiration demand, which is a cost to crops with *A*_j_ enhancement.

### Seasonal crop growth and yield dynamics

Crop cycle simulations quantify seasonal trajectories of wheat and sorghum crop attributes and generate understanding of interactions between the crop and environment (Figures 4 and S4). In this subtropical environment (Dalby, Australia) with summer-dominant rainfall, dryland cropping typically encounters late-season water stress around the time of flowering and/or during grain filling (Chenu *et al*., 2013, Hammer *et al*., 2014). While every season and situation simulated generates specific effects on the dynamics of crop growth, it is instructive to first understand interacting processes of an example season (e.g. Figures 4).

At the beginning of the crop cycle, the cumulative crop biomass increased rapidly, followed by a near-linear growth phase before growth slowed towards the end of the cycle (Figure 4a). Hence, while simulated grain mass increased after flowering it tended to plateau as growth declined during the grain-filling period. Daily biomass growth was driven by canopy photosynthesis. Canopy photosynthesis over the diurnal period was calculated on an hourly timestep by summing the instantaneous gross CO_2_ assimilation rates of the sunlit and shaded fractions, integrated over the hour, and summed over the diurnal period. Canopy photosynthesis changed dynamically over a diurnal period showing a peak mainly due to changing incoming radiation as the sun crosses the sky (Figure 4g). Over the entire crop cycle, the magnitude of the diurnal canopy photosynthesis peaks also changed dynamically and was driven by canopy LAI, SLN, and crop water status. The LAI trajectory was determined by planting density, leaf appearance and expansion rates, and leaf size. Increasing LAI increased canopy radiation interception (Figure 4b). SLN was a consequence of leaf area growth, crop N supply and demand for N by competing growing organs (Figure 4e). The drop in SLN after flowering was due to translocation of N from leaves to satisfy demands of developing grain. The SLN level determined the key photosynthetic parameters (*V*_cmax_, *V*_pmax_ and *J*_max_). Their values for the uppermost leaves of the canopy on a leaf area basis are shown for the standard temperature of 25°C (Figure 4d). As the photosynthetic parameters are temperature dependent, values calculated using the maximum temperature of the day are also shown (Figure 4d). The effect of daily temperature on canopy photosynthesis was less apparent than those of LAI and SLN. Silva-Pérez *et al*. (2017) found leaf photosynthetic rate was relatively stable across a wide range of temperature.

Canopy photosynthesis was impacted by crop water stress in the second half of the crop cycle as soil water was depleted in the exemplary crop cycle simulations. The potential demand for water uptake was driven by the transpiration rate required to maintain *C*_i_ and CO_2_ assimilation rate. If the transpiration demand could not be met by uptake and supply from the roots, then whole-crop transpiration was limited (Figure 4c). This can limit stomatal conductance, operating *C*_i_, and CO_2_ assimilation rate (e.g. Figure S1a). The severity of crop water limitation, which was indexed by the supply/demand ratio (swdef_photo) (Figure 4f), also caused leaf senescence, which reduced radiation interception (Figure 4b). The reduction in growth rate and plateau in cumulative biomass towards maturity was due to a combination of reductions in: canopy LAI, which reduced light interception; SLN, which reduced leaf and canopy photosynthetic performance; and crop water status, which reduced conductance and photosynthesis. Overall, these slowed down the grain mass/yield trajectory (Figures 4a and S4a).

The daily canopy photosynthesis peaks were influenced by the combined effects of radiation, water, temperature, LAI, and SLN (Figure 4g). Canopy photosynthesis was made up of contributions from the sunlit and shaded fractions. The shaded fraction was almost always *A*_j_ limited, while the sunlit fraction could be *A*_c_ or *A*_j_ limited. In the wheat example, the sunlit fraction was mostly *A*_j_ limited in the first half of the crop cycle (Figure 4g). However, when the crop was under water stress in the second half of the crop cycle, *A*_c_ limitation became dominant (Figure 4f,g). As explained earlier, this was due to reduced stomatal conductance and *C*_i_ (e.g. Figure S1a). The predicted *A*_c_–*A*_j_ dynamics captures the important seasonal water stress effects on canopy photosynthesis when they occur. These *A*_c_ and *A*_j_ dynamics also occurred in the sorghum example (Figure S4g). In addition, the switch between *A*_c_ and *A*_j_ limitation was more sensitive to temperature drops in sorghum. The brief dip in air temperature early in the season (Figure S4d) caused *A*_c_ limitation in the sunlit fraction (Figure S4g). The simulated sensitivity to low temperatures is consistent with C_4_ photosynthesis temperature analysis (Kubien *et al*., 2003). Such complex dynamics of crop growth and yield will unfold differently with different photosynthetic manipulations and the seasonal weather pattern.

### Crop yield response to photosynthetic manipulation is more complex than expected

Wheat and sorghum crops with and without photosynthetic manipulation were simulated across a diverse range of production environments (Figure S3 and Table S4). The simulated baseline wheat yield from the six representative sites across Australia varied widely from 0.5–6 t/ha. Dalby, Dubbo, Dookie were the higher-yielding sites (up to 6 t/ha), Katanning was in the mid-range (2–3.25 t/ha), and Walpeup and Merredin were the lower-yielding sites (0.5–3.5 t/ha, but mostly below 2.5 t/ha) (Figure 5). The variations in the baseline yield across the sites were due to local environment, agronomic management practices with N input as the major factor (Table S4), and seasonal climate variability within sites (Chenu *et al*., 2013). The simulated baseline sorghum yield from the four representative sites in NE Australia also varied widely from 1–8 t/ha. However, although agronomic management practices (Table S4) were similar, there was significant variation at all sites due to the extent of seasonal climate variability. The simulated wheat and sorghum yields in the different local environments were comparable to those reported previously in comprehensive crop-environment analysis studies (Chenu *et al*., 2013, Hammer *et al*., 2014) indicating the cross-scale model extension is robust across a spectrum of non-stressed and stressed crop conditions, as previously demonstrated (Wu *et al*., 2019).

The magnitude of yield change relative to the baseline scenario associated with photosynthetic manipulations (Δyield) was dependent on both the manipulation target and the environment (Figures 5 and 6). The top decile of Δyield was up to an equivalent of 8.1% yield increase with the installation of the full cyanobacterial-type CCM (Figure 5f). The simultaneous enhancement in Rubisco functions, Rieske-FeS protein, and mesophyll conductance gave both wheat and sorghum Δyield of up to 6.6% (Figures 5i, 6i). This yield effect was also predicted for Chlorophyll *d* and *f* in sorghum (Figure 6h). The Rubisco function (Figures 5a,b,c and 6a,b,c) and electron transport chain (Figures 5g,h, 6g) targets had similar, but smaller Δyield effects (∼1.4–3.7%) in both wheat and sorghum. Apart from reducing bundle sheath conductance in sorghum (Figure 6f), the other CO_2_ delivery related targets had limited effect on wheat and sorghum Δyield (Figures 5d and 6d,e). The comparative magnitudes of the top decile of Δyield presented in Figures 5 and 6 agreed well with the leaf-level enhancements predicted earlier (Figures 2, 3). The physiological reasons for the predicted Δyield from a whole-crop context, and its apparent variability across different environments, are discussed below.

The physiological underpinning for the predicted positive Δyield was an increased grain number in both wheat and sorghum. A detailed inspection of the predicted crop attribute trajectories revealed that leaf photosynthetic manipulations that enhanced *A*_j_ increased canopy CO_2_ assimilation and biomass growth, and canopy size (or LAI) early in the crop cycle (e.g. Figures 4 and S5). This allowed crops to achieve higher growth rates, transpiration, and biomass around anthesis, which increased grain number (van Oosterom & Hammer, 2008). In situations with positive Δyield, water and nitrogen were less limiting after anthesis, hence the crop could carry on photosynthesizing to fill all grains so grain size were not impacted (e.g. Figure S5). In these cases, enhanced leaf photosynthesis increased yield. Since the installation of the full CCM gave the largest effect on *A*_j_ (Figure 2f), it generated the largest Δyield as anticipated (Figure 5f). A large effect on rice biomass growth with a full CCM was also predicted in a previous study (Yin & Struik, 2017). Figures 2, 3 and 5, 6 show how each of the manipulation outcomes impacted yield.

However, considerable variability in Δyield was predicted even in high-yielding conditions (e.g. Figures 5f, high yield region). The physiological underpinnings of this were associated with interactions between the altered crop growth and the timing and severity of water and/or nitrogen stress around the critical flowering–grain filling period. Despite increased canopy photosynthesis and biomass growth in the first half of the crop cycle, photosynthetic enhancement caused increased transpiration and exacerbated the severity of late-season water stress in less water-abundant seasons due to higher gas-exchange rates earlier in the season (e.g. Figure S6). This resulted in reduction in stomatal conductance and CO_2_ supply for photosynthesis later in the cycle. This could be further compounded by a reduction in LAI due to enhanced leaf senescence reducing canopy light interception. Greater early biomass growth increases crop N demand and generates a later dilution of leaf nitrogen causing lower SLN and photosynthesis in the second half of the crop cycle. The overall result would be lower canopy photosynthesis and crop growth rates during the grain-filling period, resulting in reduced grain size. In some instances, such grain size reduction would offset grain number increase, thus explaining the Δyield variability (e.g. Figures 5g,h and 6g,h). This highlights the fact that effects of photosynthetic enhancement will be modulated by whole-plant physiological limits and the environmental context especially in resource (water and nitrogen) limited production environments.

The nature of Δyield and its variability in high-yielding conditions differed between the manipulation targets. Manipulations that enhanced *A*_c_, including Rubisco function and the full CCM, resulted in Δyield that ranged from near nil to small positive values (Figures 5a,b,c,f and 6a,b,c). However, some negative Δyield were predicted with manipulations that enhanced *A*_j_, including the electron transport chain targets (Figures 5g,h and 6g,h). The manipulation target stacking scenario resulted in wider Δyield variations than its component targets (Figures 5i and 6i). Given the consequence of the manipulation on timing and severity of water and/or nitrogen stress, Rubisco functions, the installation of the full cyanobacterial-type CCM, or reduced bundle sheath conductance manipulations (Figures 5c,f and 6f) should also result in negative Δyield outcomes as with the electron transport chain targets (Figures 5g,h and 6g,h). However, this was predominantly not the case due to the benefit of *A*_c_ enhancement in improving canopy photosynthesis especially under water stress conditions. The Rubisco and CO_2_ delivery targets resulted in improved canopy photosynthesis and biomass growth during the stress period through better intrinsic water use efficiency (e.g. Figure S1a). This means enhanced canopy photosynthesis, crop growth, and less impacts on grain size. As expected, the manipulation target stacking scenario slightly improved the negative Δyield compared with enhancing the electron transport chain targets (Figures 5i and 6i).

Variability of Δyield in the low-yielding conditions was also dominated by the timing and severity of water and/or nitrogen stress as for the high-yielding conditions. These were characterized by the 10^th^ and 90^th^ percentile regressions (Figures 5 and 6). The regressions also highlighted that manipulation strategies generating enhanced *A*_j_ were especially beneficial for the high-yielding environments as there were instances that Δyield increased well above the general trends (Figures 5g,h,i and 6g,h,i). This was due to *A*_j_ being the predominant limitation over the crop cycle (e.g. Figures 4g and S4g) and in seasons where more water was available, increased crop water use was less detrimental. Although there are modest gains to be made with the best photosynthetic manipulation strategies (e.g. Figures 5f and 6h,i), another key for crop improvement is better addressing the variation in Δyield generated by plant–environment interactions.

Installation of the cyanobacterial *HCO* transporters showed a distinct Δyield pattern (Figure 5e). The positive Δyield was not due to increased grain number as described earlier. Analysis revealed that canopy photosynthesis, biomass growth, and grain number were reduced (e.g. Figure S7). Canopy photosynthesis was reduced early in the crop cycle due to the extra ATP costs of the transporters reducing the already limiting *A*_j_ (Figure 4g). The decline in canopy-level *A*_j_ was consistent with the leaf-level result (Figure 2e). However, reduced *A*_j_ and growth helped conserve water and nitrogen for the second half of the crop cycle. This meant better LAI retention, canopy light interception, and water availability, so growth rates were better sustained during the critical flowering–grain filling period and increased grain size, which compensated for the reduction in grain number due to reduced early seasons growth. However, the *HCO* transporters installation was also the only approach that resulted in large negative Δyield effects (Figure 5e). In contrast to the negative Δyield with some of the other manipulation cases (e.g. Figure 5g,h), this occurred in those seasons with more plentiful water and nitrogen conditions where grain size was close to its potential so any reduction in grain number led to sink limitation and reduced yield.

### Case study: Potential impact on Australian crop production and globally

Quantifying the potential impact of leaf photosynthetic manipulation strategies on Australian wheat and sorghum production at national scale revealed differences among the manipulation targets and crops. Potential magnitude of enhancement in the predicted steady-state *A*_c_ and *A*_j_ (Figures 2, 3) reflected expectations based on transgenic and modelling studies. However, the largest levels of crop production increase were modest with median increases of 3–4% at national scale (Figure 7). The modest levels of increase, which exhibit a range of potential outcomes and instances of negative change, were associated with more rigorous sampling of effects of diverse environmental and agronomic conditions that generate a realistic frequency of incidence of water and nitrogen limitations at national scale. The full CCM installation (4.2) generated the largest increase in Australian wheat production with a median gain of ∼3%, while some of the Rubisco (1.1 and 2) and bicarbonate transporter (4.1) manipulation strategies generated ∼1% increase. Rieske Fe-S (7) and chlorophyll *d* & *f* (8) manipulation strategies resulted in slightly reduced overall production at the national scale. The electron transport chain targets resulted in wider production change variabilities as they tended to exacerbate crop water and/or nitrogen stress. The manipulation stacking strategy (9) did not result in further increase in the median value compared to just “better” Rubisco (2), but it also increased the production variability. In sorghum, incorporating chlorophyll *d* and *f*, and the manipulation stacking strategy generated the largest production gain (3–4%). This contrasted with the wheat predictions as nitrogen limitation was less detrimental in sorghum production. Nitrogen deficiency was also found to reduce yield gains with enhanced photosynthesis from elevated CO_2_ in a large number of C_3_ crops (Ainsworth & Long, 2021). Other manipulation targets such as those related to Rubisco (1.3, 1.4, and 2), bundle sheath conductance (6), and Rieske Fe-S (7) will likely result in ∼1–2% increase in Australian sorghum production. In both crops, the likely impact of manipulating mesophyll conductance (3) was consistently low.

**Figure 7.**
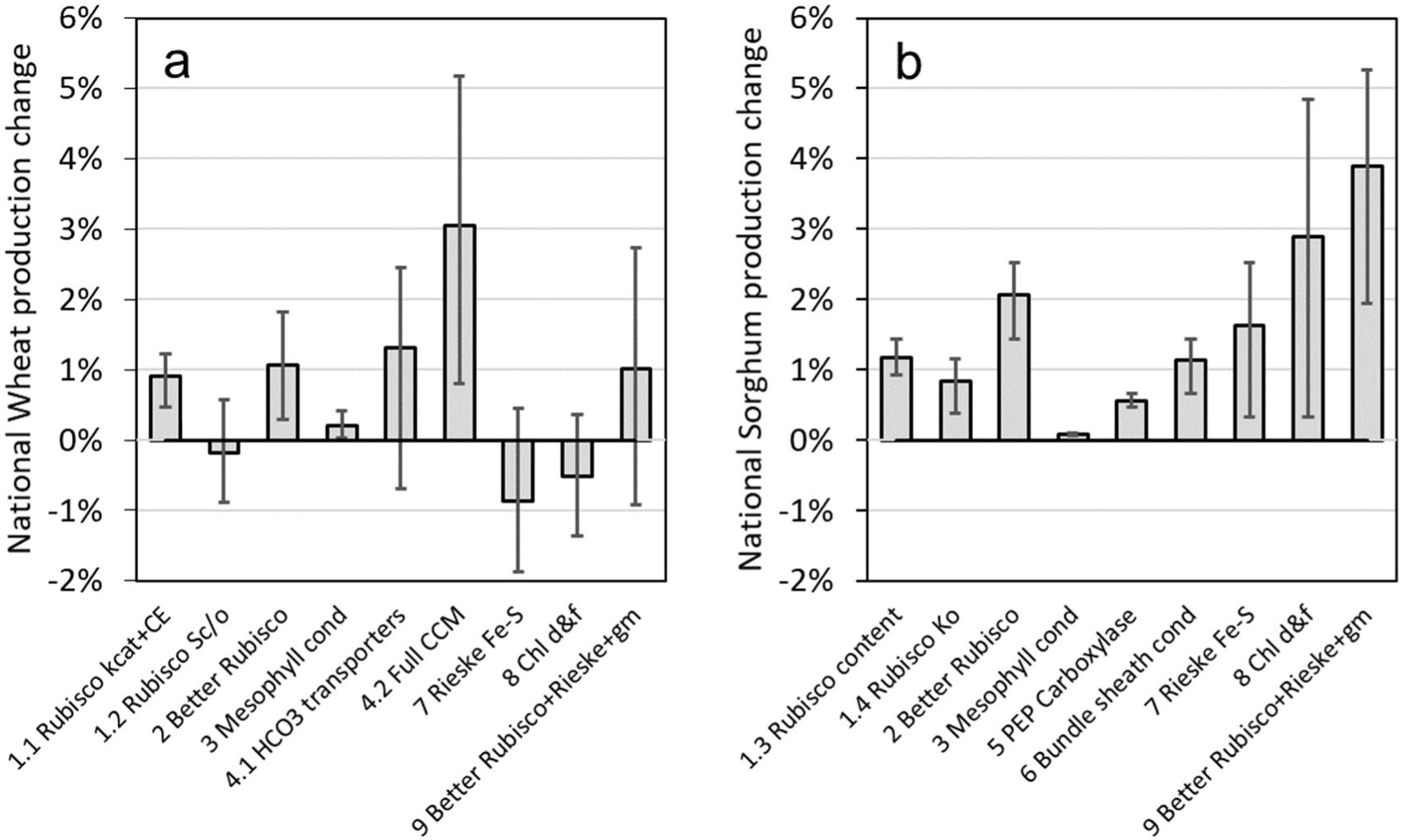
Predicted percentage change in Australia-wide (a) wheat and (b) sorghum production associated with leaf photosynthetic manipulations (Table 1). This expanded set of simulation uses representative seasonal weather data sampled from the past 120 years, three representative levels of each of sowing date and starting soil water specific for each site (Table S4). Median values are given by bars. Whiskers show the first and third quartile values, which are calculated using the corresponding quartile values from all production sites (wheat: n = 12,960 crop cycles per bar; sorghum: n = 8,640 crop cycles per bar).

Benchmarking impacts of photosynthetic enhancement against current year-on-year crop yield advances provides a useful context for crop breeding efforts. The historical annual rates of increase in the national average yield of Australian wheat and sorghum are 1.2% and 2.1% respectively (Potgieter *et al*., 2016). These rates quantify the extent of continual technological advances arising from crop improvement due to empirical breeding based on selection for yield, advances in agronomy such as stubble management practices to enhance soil water availability, and some environmental trend effects (e.g. rising CO_2_). Hence, implementing the best of the photosynthetic manipulation targets will likely result in an equivalent of 2.5 and 2 years of conventional production gains for Australian wheat and sorghum, respectively.

An effective first-order approach in predicting yield impacts across international locations is by applying the predicted Australian yield changes to correlated international environments globally. Australian production environments present a broad spectrum of non-stressed to stressed production conditions. Marginal Australian environments like southern and western Australia are correlated with South America, southern Africa, Iran and high latitude European and Canadian locations (Mathews *et al*., 2007). For these locations, the installation of full CCM is likely to be the most beneficial, generating between 0.1% and up to 8.1% gains based on the top and bottom 10^th^ percentile regression (Figure 5f). High-yielding environments such as eastern Australia are correlated with international locations including the Indo-Gangetic plains, West Asia, North Africa, Mexico, and locations in Europe and Canada (Mathews *et al*., 2007). In these environments and if water and nitrogen were also abundant, the manipulation stacking strategy is likely to be the most beneficial, generating up to 15% gains in wheat yield (Figure S8). This is also evident in Figure 5i showing instances of large Δyield well above the top 10th percentile regression. Understanding and quantifying production environment context dependencies is important for maximizing yield improvement.

### Cross-scale analysis helps understand and quantify effects on crop yield

This study used a state-of-the-art cross-scale model to predict effects of a broad list of photosynthetic manipulation strategies on seasonal crop growth and yield dynamics and quantified the potential impact (or lack of it) on crop yield across a broad spectrum of non-stressed to stressed production conditions. Based on the potential magnitude of enhancement in the steady-state leaf photosynthetic rates, predicted yield increases are likely modest even in the top decile of seasonal outcomes. Importantly, yield change can vary from the top seasonal outcomes, which ranging between 0 and 8% depending on the crop type and manipulation, to nil or losses depending on availability of water and nitrogen. The modest results and environmental context dependencies challenge common perceptions, which have been based on limited field experiments and modelling, of the magnitude of benefits likely to arise from photosynthetic manipulation. Our analysis on the manipulation of the steady-state enzyme- and electron transport-limited photosynthetic rates suggests a multi-pronged approach to enhance both will be needed for achieving larger yield gains, which will likely be achieved by stacking Rubisco function and electron transport chain enhancements or installing a full CO_2_ concentrating system. There is also a need to address environmental context dependencies confounding yield improvement to maximise yield impact from the photosynthetic manipulations. With models that connect processes and interactions across biological scales of organization, integrated systems analysis from leaf to yield can be used to assess multitudes of potential photosynthetic targets (Zhu *et al*., 2020) and generate novel information to help accelerate photosynthesis research and crop improvement (Chew *et al*., 2017, Hammer *et al*., 2019).

## Supporting information

Supplemental Equations, Tables, Figures

## Acknowledgements

This research was funded by grants from the Australian Research Council: Centre of Excellence for Translational Photosynthesis CE1401000015 (All) and DE210100854 (A.W.). We thank Prof. Mark Cooper for advice on applying results from this study to international wheat production environments.

## Supplemental Material

Eqns 1–20: Expanded photosynthesis–CO_2_ diffusion model equations

Figure S1. Wheat sunlit

Figure S2. Sorghum sunlit

Figure S3. Australia crop production regions and key sites in each of the region.

Figure S4. Predicted sorghum crop attributes dynamics, and environmental variables over a sample crop cycle.

Figure S5. Wheat crop attributes trajectories with abundant water and N (baseline vs target 4.2).

Figure S6. Wheat crop attributes trajectories with limited water (baseline vs target 4.2).

Figure S7. Wheat crop attributes trajectories with reduced growth (baseline vs target 4.1).

Figure S8. Dalby wheat and sorghum +I+N (300kg/ha total), mid sowing, 9 bars 9 manipulations. % yield change.

Figure S9. CCM with less *V*_bmax_ (PsiVp=0.41)

Table S1. CO_2_ diffusion parameters of the photosynthesis–CO_2_ diffusion model

Table S2. Lumped coefficients of the photosynthesis–CO_2_ diffusion model

Table S3. Parameters associated with electron transport

Table S4. Crop production sites, cultivars, and agronomic practices used in simulations.

Table S5. Photosynthesis model parameters and their baseline values.

Table S6. Long-term average crop production data from the Australian Bureau of Statistics.

Table S7. Predicted regional production change percentage.

## Notes

**Funding:** This research was supported by grants from the Australian Research Council: Centre of Excellence for Translational Photosynthesis CE1401000015 (All) and DE210100854 (A.W.).

### Competing Interest Statement

The authors have declared no competing interest.

