## Supplemental Equations, Tables, Figures for "A cross-scale analysis to understand and quantify effects of photosynthetic enhancement on crop growth and yield"

### Appendix A: Expanded photosynthesis–$CO_2$ diffusion model equations

Three photosynthetic pathways are modelled: $C_3$, $C_4$, and single-cell CCM. Equations for calculating the Rubisco-limited ($A_c$) and electron-transport-limited ($\underline{A}_j$) rates of $CO_2$ assimilation, and the chloroplastic $CO_2$ supply function:

The $C_3$ model (Farquhar *et al.*, 1980):

$$A_c = \frac{(C_c - \Gamma_*)V_{cmax}}{C_c + K_c(1 + O_c/K_o)} - R_d \qquad \text{Eqn 1}$$

$$A_j = \frac{(C_c - \Gamma_*)J}{4C_c + 8\Gamma_*} - R_d \qquad \text{Eqn 2}$$

$$C_c = C_i/C_a \times C_a - \frac{A}{g_m} \qquad \text{Eqn 3}$$

where $V_{cmax}$ is the maximum Rubisco activity; $K_c$ and $K_o$ are the Rubisco Michaelis-Menten constants for $CO_2$ and $O_2$ respectively; $C_a$ and $C_i$ are the ambient and intercellular $CO_2$ partial pressures, and $C_c$ and $O_c$ are the chloroplastic $CO_2$ and $O_2$ partial pressures; $\Gamma_*$ is the $CO_2$ compensation partial pressure in the absence of mitochondrial respiration ($R_d$) as calculated by half $O_c$ multiplied by the reciprocal of the relative specificity of Rubisco (i.e. $0.5O_c/S_{c/o}$); $J$ is potential electron transport rate and is related to the incident irradiance as in Eqn 4; $g_m$ is the mesophyll conductance to $CO_2$. The $CO_2$ supply function uses a $C_i/C_a$ ratio and the Fick's first law of diffusion.

The relationship between $J$ and irradiance ($I$) is modelled by the solution to a non-rectangular hyperbolic-type function (Farquhar & Wong, 1984, von Caemmerer, 2000):

$$J = \frac{\alpha_{PSII}I + J_{max} - \sqrt{(\alpha_{PSII}I + J_{nax})^2 - 4\theta\alpha_{PSII}IJ_{max}}}{2\theta} \qquad \text{Eqn 4}$$

Where $J_{max}$ is maximum electron transport and $\theta$ is an empirical curvature factor of the *J-I* response. $\alpha_{PSII}$ is the fraction of *I* absorbed and used by Photosystem II to drive electron transport. $\alpha_{PSII}$ is calculated by assuming that leaves absorb 85% of the incident *I*, of which 15% is not useful due to spectral quality, of which half goes to Photosystem II (i.e. 0.85 * (1 – 0.85) * 0.5 = 0.36) (von Caemmerer, 2000). Compared with the widely used relationship between *J* and the quantum yield of Photosystem II from chlorophyll fluorescence measurement (Maxwell & Johnson, 2000), Eqn 4 is used to describe *J–I* over a range of irradiance here.

The $C_4$ model (von Caemmerer, 2000):

$$A_\mathrm{c} = \frac{(C_\mathrm{s} - \gamma_* O_\mathrm{s}) V_\mathrm{cmax}}{C_\mathrm{s} + K_\mathrm{c}(1 + O_\mathrm{s}/K_\mathrm{o})} - R_\mathrm{d} \qquad \text{Eqn 5}$$

$$A_\mathrm{j} = \frac{(C_\mathrm{s} - \gamma_* O_\mathrm{s})(1-x) J_\mathrm{t}}{3C_\mathrm{c} + 7\gamma_* O_\mathrm{s}} - R_\mathrm{d} \qquad \text{Eqn 6}$$

$$C_\mathrm{s} = C_\mathrm{i}/C_\mathrm{a} \times C_\mathrm{a} - \frac{A}{g_\mathrm{m}} + \frac{P - A - R_\mathrm{m}}{g_\mathrm{bs}} \qquad \text{Eqn 7}$$

where $C_\mathrm{s}$ and $O_\mathrm{s}$ are the bundle-sheath chloroplastic $CO_2$ and $O_2$ partial pressures; $\gamma_*$ is the reciprocal of the relative specificity of Rubisco; here $J_\mathrm{t}$ is to indicate that electron transport is modelled as a whole and allocated the $C_3$ and $C_4$ cycles; $x$ is the fraction of the generated ATP required in the mesophyll; $R_\mathrm{m}$ is the respiration occurring in the mesophyll; $g_\mathrm{bs}$ is the bundle-sheath conductance to $CO_2$. This $CO_2$ supply function for $C_4$ photosynthesis is based on the $C_3$ version with an added active transport of $CO_2$ into the relative air-tight bundle sheath involving initially fixing of $CO_2$ by PEP carboxylase in the mesophyll. Due to the air-tight nature, evolved of $O_2$ would elevate $O_\mathrm{s}$, which is given by Eqn 9. $P$ is the $CO_2$ supply from the mesophyll that depends on catalytic properties of PEP carboxylase and the limitation state of $CO_2$ assimilation (Eqn 8).

PEP carboxylation rate is defined, using equations from von Caemmerer (2000), by:

$$P = \begin{cases} \min\left(\frac{C_\mathrm{m} V_\mathrm{pmax}}{C_\mathrm{m} + K_\mathrm{p}}, V_\mathrm{pr}\right) & \text{, for } A_\mathrm{c} \\ xJ/\varphi & \text{, for } A_\mathrm{j} \end{cases} \qquad \text{Eqn 8}$$

Where $C_\mathrm{m}$ is the mesophyll $CO_2$ partial pressure; $V_\mathrm{pmax}$ is the maximum PEP carboxylase activity and $K_\mathrm{p}$ is the PEP carboxylase Michaelis-Menten constants for $CO_2$; $\varphi$ is the 2 ATP required by the $C_4$ cycle.

Because of the relative air-tight nature of the bundle sheath, $O_2$ evolved in the cell will increase its partial pressure beyond the mesophyll $O_2$ partial pressure ($O_\mathrm{m}$). This is calculated by Farquhar (1983) and von Caemmerer (2000):

$$O_\mathrm{s} = O_\mathrm{m} + \frac{\alpha A}{0.047 g_\mathrm{s}} \qquad \text{Eqn 9}$$

Where the second term gives the extra $O_2$ proportional to $A$ and the fraction of $O_2$ evolution occurring in the bundle sheath ($\alpha$).

The single-cell CCM model is derived from the $C_4$ and $C_3$ models as described by Price *et al.* (2011) and von Caemmerer (2021):

$$A_c = \frac{(C_x - \gamma_* O_x) V_{cmax}}{C_x + K_c(1 + O_x/K_o)} - R_d \qquad \text{Eqn 10}$$

$$A_j = \frac{(C_x - \gamma_* O_x) z(1-x) J}{3C_x + 7\gamma_* O_x} - R_d \qquad \text{Eqn 11}$$

$$C_x = C_i/C_a \times C_a - \frac{A}{g_w} + \frac{P - V_c}{g_x} \qquad \text{Eqn 12}$$

where $C_x$ and $O_x$ are the carboxysomal $CO_2$ and $O_2$ partial pressures. $O_x$ is calculated using Eqn 9. Here the ATP-limited version of $A_j$ is used and $x$ is the fraction of the produced ATP required by bicarbonate pumps involved in the active transport of inorganic carbon from the mesophyll cytosol into the chloroplast. This extra ATP requirement is met by additional cyclic electron flow in the electron transport chain (Yin & Struik, 2012). The overall ATP production is calculated based on scaling the yield from linear electron flow with a factor ($z$) that captures the additional contribution from cyclic electron flow and proton requirement per ATP generated (Eqn 13). The $CO_2$ supply term into the chloroplast differs from the standard formulation of the $C_4$ model, assuming that photorespiratory $CO_2$ loss occurs in the mesophyll cytosol (Price *et al.*, 2011). $P$ is the $CO_2$ supply from the mesophyll cytosol that depends on catalytic properties of the bicarbonate transporters and the limitation state of $CO_2$ assimilation (Eqn 14) and $V_c$ is the Rubisco carboxylation rate (Eqn 15); $g_w$ and $g_x$ are the mesophyll cell wall/plasmalemma and the combined carboxysome–chloroplast envelope conductance to $CO_2$ respectively.

The rate of ATP production with additional contribution from cyclic electron flow is given by von Caemmerer (2021):

$$J_{ATP} = \frac{3 - f_{cyc}}{h(1 - f_{cyc})} J = zJ \qquad \text{Eqn 13}$$

where $f_{cyc}$ is the fraction of the electron flow out of Photosystem I that procedes via cyclic electron flow; $h$ is the number of protons required per ATP generated, which is taken to be 4.

The rate of bicarbonate transporters is given by Price *et al.* (2011):

$$P = \begin{cases} \frac{C_{cy}\ V_{bmax}}{C'_{cyt} + K_b} & \text{, for } A_c \\ zxJt/\varphi & \text{, for } A_j \end{cases} \qquad \text{Eqn 14}$$

Where $C_{cyt}$ is the mesophyll cytosol $CO_2$ partial pressure; $V_{bmax}$ is the maximum bicarbonate transporter activity and $K_p$ is the bicarbonate transporter Michaelis-Menten constants for $CO_2$.

$\varphi$ is the ATP requirement of the two types of cyanobacterial bicarbonate transporter (i.e. BicA and SbtA; requiring 0.25 and 0.5 ATP, respectively, per bicarbonate transported) (Price *et al.*, 2011).

The Rubisco carboxylation rate ($V_c$) is given by either the RuBP-saturated carboxylation rate or the possible rate supported by the electron transport:

$$V_c = \begin{cases} \dfrac{C'_x V_{cmax}}{C'_x + K_c(1 + O'_x/K_o)} & \text{, for } A_c \\ \dfrac{C'_x z(1-x)J}{3{C_x}' + 7\gamma_* O'_x} & \text{, for } A_j \end{cases} \qquad \text{Eqn 15}$$

The expanded equation of the three photosynthesis models can be generalised into a single equation by combining Eqns 1–15:

$$A_\xi = \frac{\left\{\left((p - qA_\xi) + \dfrac{\left[\left((p-qA_\xi)x_4 + x_5\right) - x_6 A_\xi - R_m - x_7\right]x_8}{s}\right) - (x_{10}A_\xi + x_{11})\right\} x_1}{\left((p - qA_\xi) + \dfrac{\left[\left((p-qA_\xi)x_4 + x_5\right) - x_6 A_\xi - R_m - x_7\right]x_8}{s}\right) + x_{12}A_\xi + x_{13} + x_3} - R_d \qquad \text{Eqn 16}$$

Where $CO_2$ diffusion variables are given in Table S1. The $x$'s are lumped coefficients given in Table S2.

Table S1. Variables associated with gas ($CO_2$ and $O_2$) diffusion used in Eqn 16.

| Stomatal mode | $p$ | $q$ | $b$ | $s$ |
|---|---|---|---|---|
| Non-limiting water: constant $C_i/C_a$ | $C_i/C_a \times C_a$ | $\dfrac{1}{g_m}$ | $\dfrac{\alpha}{0.047}$ | $g_{bs}$ or $g_x$ |
| Water limited: conductance from water supply-demand balance | $C_a - \dfrac{TC_a}{g_t + T/2}$ | $\dfrac{1}{g_t + T/2} + \dfrac{1}{g_m}$ | $\dfrac{\alpha}{0.047}$ | $g_{bs}$ or $g_x$ |

Table S2. Lumped coefficients used in Eqn 16 for the different photosynthesis models. Electron transport associated parameters (i.e. $x$, $z$ and $\varphi$) are defined in Table S3.

| | $x_1$ | $x_3$ | $x_4$ | $x_5$ | $x_6$ | $x_7$ | $x_8$ | $x_{10}$ | $x_{11}$ | $x_{12}$ | $x_{13}$ |
|---|---|---|---|---|---|---|---|---|---|---|---|
| | $C_3$ model | | | | | | | | | | |
| $A_c$ | $V_{cmax}$ | $K_c$ | 0 | 0 | 0 | 0 | 0 | 0 | $\gamma_* O_m$ | 0 | $\frac{K_c}{K_o} O_m$ |
| $A_j$ | $J/4$ | 0 | 0 | 0 | 0 | 0 | 0 | 0 | $\gamma_* O_m$ | 0 | $2\gamma_* O_m$ |
| | $C_4$ model | | | | | | | | | | |
| $A_{c1}$ (PEP-saturated rate) | $V_{cmax}$ | $K_c$ | $V_{pmax}/(C_m' + K_p)$ | 0 | 1 | 0 | 1 | $\frac{\gamma_* \alpha}{0.047 g_{bs}}$ | $\gamma_* O_m$ | $\frac{K_c}{K_o} \cdot \frac{\alpha}{0.047 g_{bs}}$ | $\frac{K_c}{K_o} O_m$ |
| $A_{c2}$ (PEP-regeneration-limited rate) | $V_{cmax}$ | $K_c$ | 0 | $V_{pr}$ | 1 | 0 | 1 | $\frac{\gamma_* \alpha}{0.047 g_{bs}}$ | $\gamma_* O_m$ | $\frac{K_c}{K_o} \cdot \frac{\alpha}{0.047 g_{bs}}$ | $\frac{K_c}{K_o} O_m$ |
| $A_j$ | $(1-x)J_t/3$ | 0 | 0 | $xJ_t/\varphi$ | 1 | 0 | 1 | $\frac{\gamma_* \alpha}{0.047 g_{bs}}$ | $\gamma_* O_m$ | $\frac{7}{3} \cdot \frac{\gamma_* \alpha}{0.047 g_{bs}}$ | $\frac{7}{3} \gamma_* O_m$ |
| | Single-cell CCM model | | | | | | | | | | |
| $A_c$ | $V_{cmax}$ | $K_c$ | $V_{pmax}/(C_m' + K_p)$ | 0 | 0 | $\frac{C_x' V_{cmax}}{C_x' + K_c(1 + O_x'/K_o)}$ | 1 | $\frac{\gamma_* \alpha}{0.047 g_x}$ | $\gamma_* O_m$ | $\frac{K_c}{K_o} \cdot \frac{\alpha}{0.047 g_x}$ | $\frac{K_c}{K_o} O_m$ |
| $A_j$ | $z(1-x)J_t/3$ | 0 | 0 | $zxJ_t/\varphi$ | 0 | $\frac{C_c' z(1-x)J}{3C_x{}' + 7\gamma_* O_x'}$ | 1 | $\frac{\gamma_* \alpha}{0.047 g_x}$ | $\gamma_* O_m$ | $\frac{7}{3} \cdot \frac{\gamma_* \alpha}{0.047 g_x}$ | $\frac{7}{3} \gamma_* O_m$ |

Table S3. Parameter associated with electron transport as in Table S2.

| Model | $\varphi$ | $f_{cyc}$ | $x$ | $z$ |
|---|---|---|---|---|
| $C_4$ | 2 | $0.25\varphi$ | $\frac{\varphi}{3+\varphi}$ | $\frac{3 - f_{cyc}}{4(1 - f_{cyc})}$ |
| Single-cell CCM | 0.75 | | | |

Eqn 16 can be rearranged into a quadratic equation and $A_\xi$ can be solved to give the following general solution for the Rubisco-limited ($A_c$) and electron-transport-limited ($A_j$) rates of $CO_2$ assimilation for the $C_3$, $C_4$, and single-cell CCM pathways:

$$A_\xi = \frac{-\sqrt{a^2-4b}+a}{2d} \qquad \text{Eqn 17}$$

where

$$a = p\,s - q\,R_\mathrm{d}\,s + q\,s\,x_1 + s\,x_1\,x_{10} + R_\mathrm{d}\,s\,x_{12} + s\,x_{13} + s\,x_3 - R_\mathrm{m}\,x_8 + p\,x_4\,x_8 - q\,R_\mathrm{d}\,x_4\,x_8 + q\,x_1\,x_4\,x_8 + x_5\,x_8 - R_\mathrm{d}\,x_6\,x_8 + x_1\,x_6\,x_8 - x_7\,x_8 \qquad \text{Eqn 18}$$

$$b = (-q\,s - q\,x_4\,x_8 + s\,x_{12} - x_6\,x_8)(-R_\mathrm{m}\,R_d\,x_8 + R_\mathrm{m}\,x_1\,x_8 + p\,R_\mathrm{d}\,s + p\,R_\mathrm{d}\,x_4\,x_8 - p\,s\,x_1 - p\,x_1\,x_4\,x_8 + R_\mathrm{d}\,s\,x_{13} + R_\mathrm{d}\,s\,x_3 + R_\mathrm{d}\,x_5\,x_8 - R_\mathrm{d}\,x_7\,x_8 + s\,x_1\,x_{11} - x_1\,x_5\,x_8 + x_1\,x_7\,x_8) \qquad \text{Eqn 19}$$

$$d = q\,s - s\,x_{12} + q\,x_4\,x_8 + x_6\,x_8 \qquad \text{Eqn 20}$$

### Appendix B: Supporting Figures and Tables

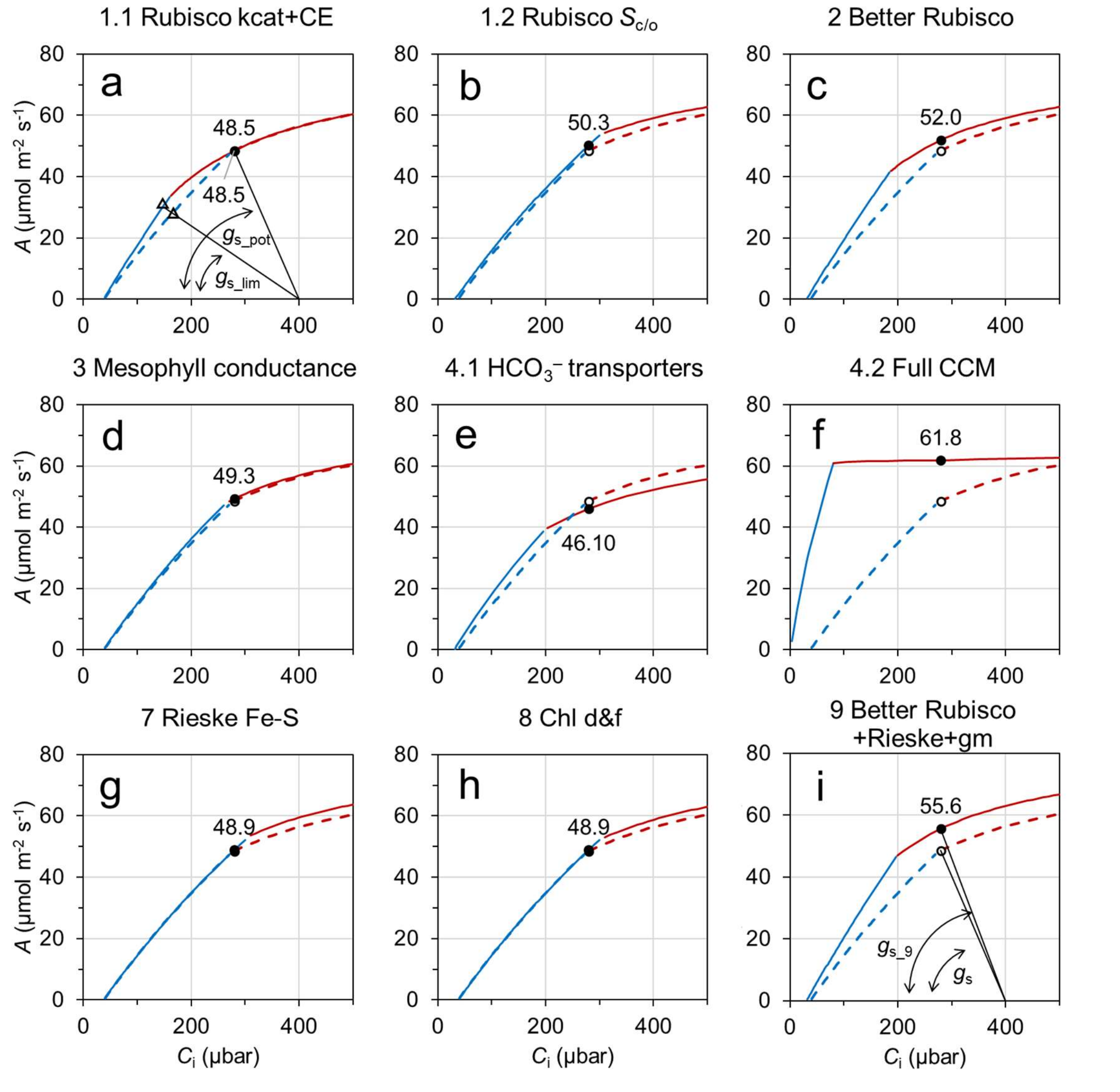


Figure S1. Simulated $C_3$ wheat sunlit fraction $A$–$C_i$ with and without photosynthetic manipulations. The manipulations are detailed in Table 1 of the main text. Calculation of the $A_c$ and $A_j$ curves, on a ground area basis is described in the main text. Slopes relating air $CO_2$, $C_i$, $A$ and $g_s$ can be drawn: e.g. (a) shows $A$ and $g_s$ in non-water limited ($g_{s\_pot}$), using a $C_i/C_a$ of 0.7, and a water limited ($g_{s\_lim}$) cases. Without affecting $g_s$, $A_c$ enhancement increases $A$, but further reduces $C_i/C_a$ under water limitation; (i) shows increase in $g_s$ with enhancement in $A_j$, using a $C_i/C_a$ of 0.7. Similarly, changes in $C_i$, $A$ and $g_s$ can be determined in other panels with and without water limitation. Results for the shaded fraction are dominated by the $A_j$ limitation across a wide range of $C_i$ due to low light levels (data not shown).

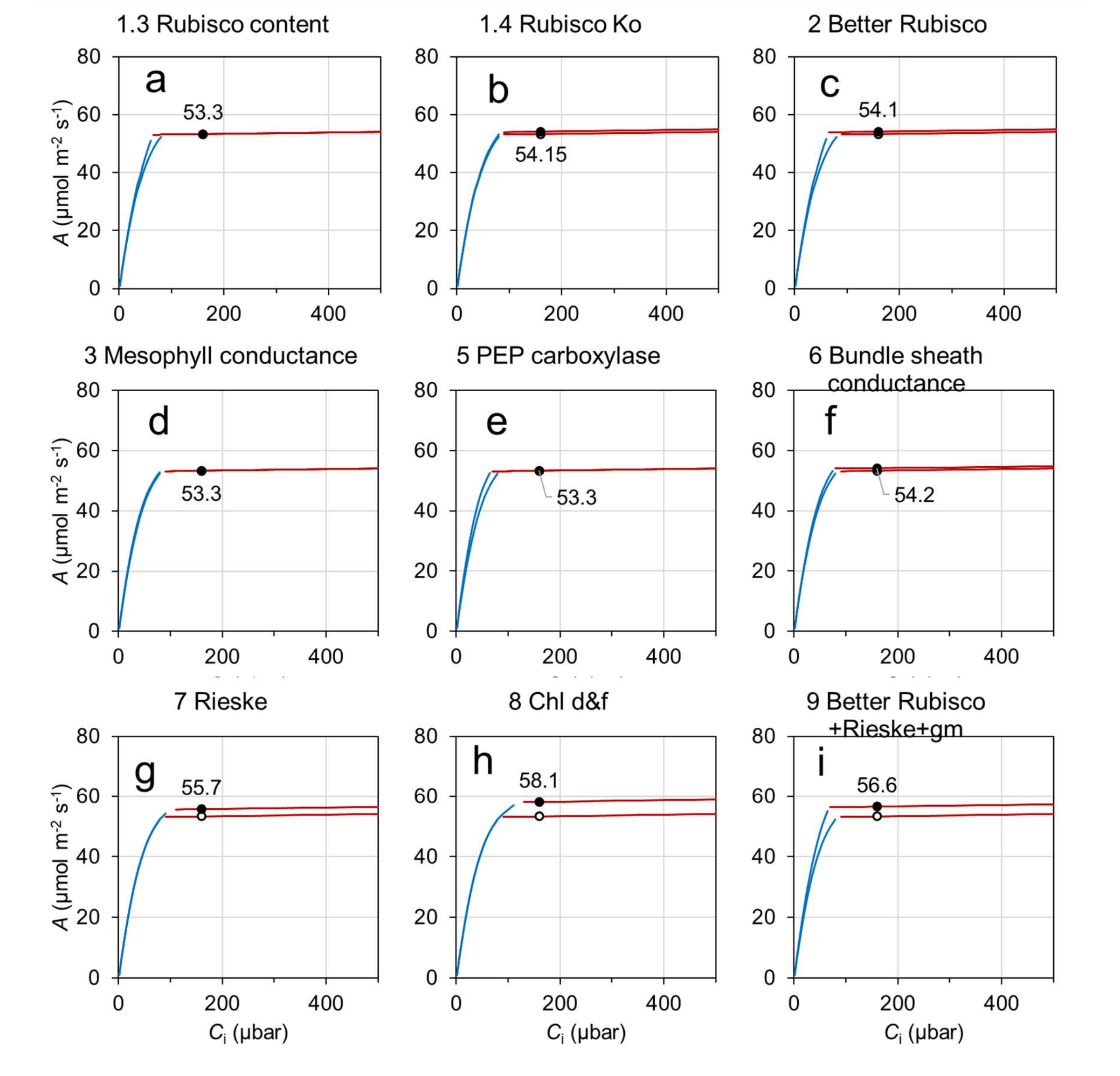


Figure S2. Simulated $C_4$ sorghum sunlit leaf fraction $A$–$C_i$ with and without photosynthetic manipulations. The manipulations and $A_c$, $A_j$ calculations are detailed in the main text. A $C_i/C_a$ of 0.4 is used for sorghum. Figure descriptions are analogous to that of Figure S1.

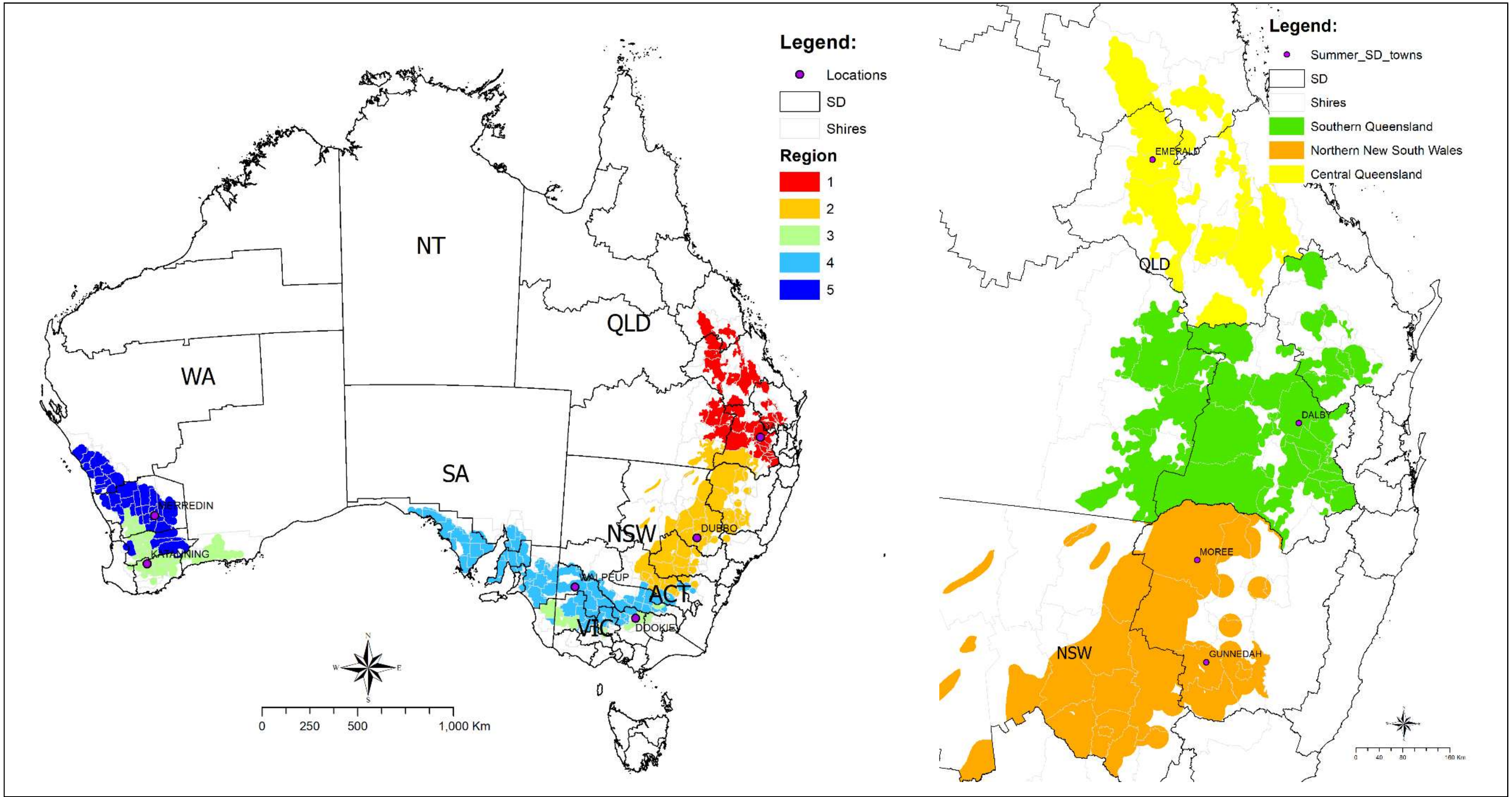


Figure S3. Australian wheat (left) and sorghum (right) production regions and the key site(s) in each region (Table S4). Statistical division (SD) boundaries are depicted in solid black lines (www.abs.gov.au).

157

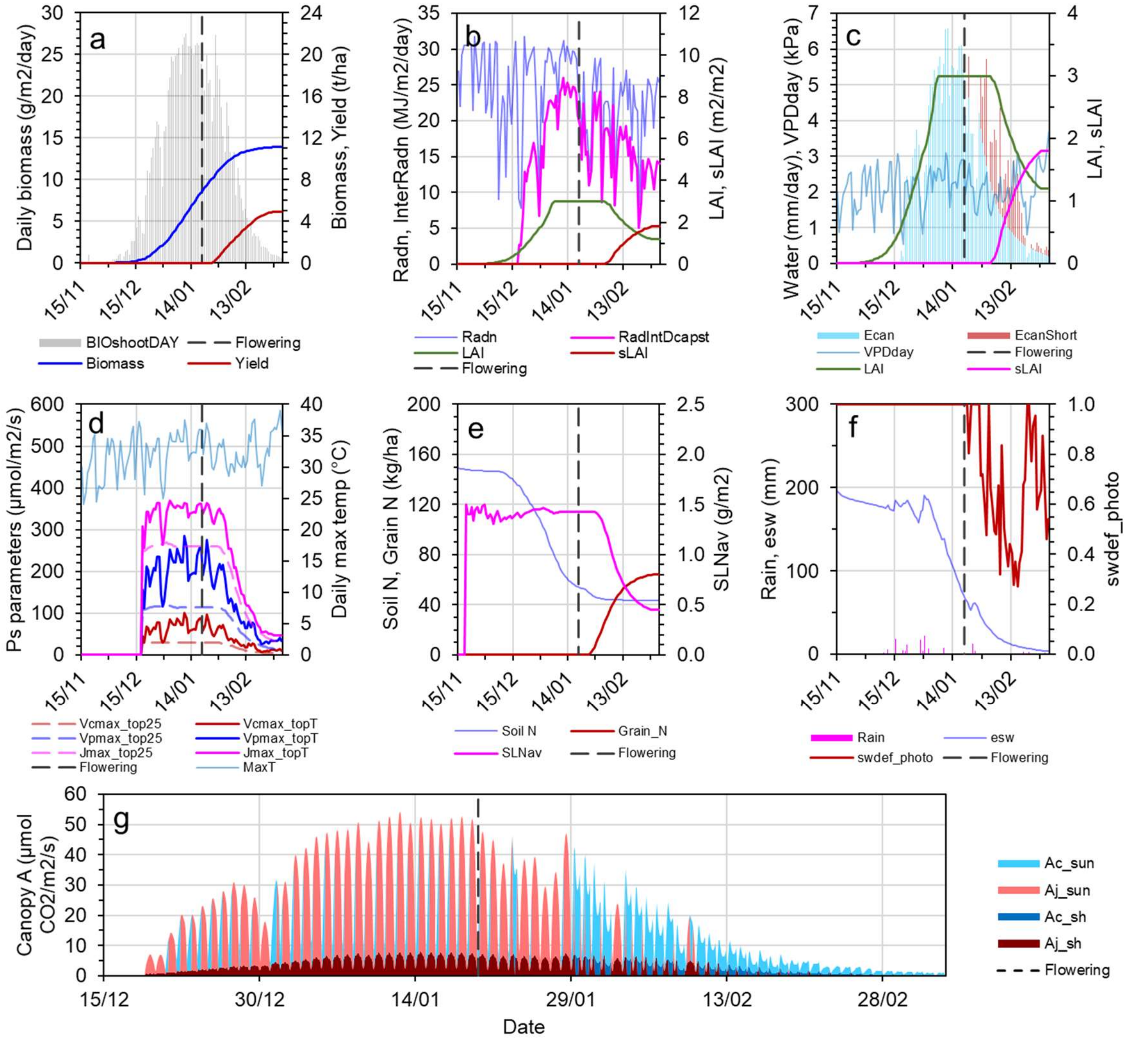


Figure S4. Predicted sorghum crop attributes dynamics, and environmental variables over a sample crop cycle. Results are from a medium-yielding year at the Dalby site with the medium sowing date and starting soil water (Table S4). (a) Cumulative crop biomass and yield. (b) Canopy leaf area index, solar radiation and interception. (c) Potential crop water demand is shown by the bars, which is made up of a fraction that is met by supply from soil water uptake by roots (i.e. actual water use) and a fraction that is not met (red bars). (d) Photosynthetic parameters for the uppermost leaves of the canopy at 25°C and the maximum air temperature during the day. (e) Soil N supply and crop N status including specific leaf nitrogen and N in grains. (f) Plant extractable soil water and a crop water stress factor; a value of 1 means all crop water demand is being met, while 0 means no water is available. (g) Daily canopy photosynthesis; each peak is made up of a histogram of total canopy photosynthesis on an hourly timestep over one diurnal period. Abbreviations: $BIO_{shootDAY}$, daily shoot biomass growth; Radn, daily incident solar radiation, RadIntDcapst, daily intercepted radiation by the whole canopy; LAI, leaf area index; sLAI, senescenced LAI;

Ecan, actual crop water use; EcanShort, fraction of the potential demand not met by supply; VPDday, indicative daytime vapour pressure deficit; Vcmax_top25, Vpmax_top25, Jmax_top25 are the values of the maximum rate of Rubisco carboxylation, maximum rate of PEP carboxylation, and maximum rate of electron transport at infinite light at 25°C; Vcmax_topT, Vpmax_topT, Jmax_topT are those photosynthetic parameter values calculated using the maximum temperature of the day (MaxT); SLNav, canopy-average specific leaf nitrogen; esw, plant extractable soil water; swdef_photo, a crop water stress factor given by EcanFilled divided by the sum of EcanFilled and ECanShort; Ac_sun and Aj_sun, Rubisco activity and electron transport limited gross $CO_2$ assimilation rate of the sunlit leaf fraction of the canopy (only the lower of the two limitations is shown); Ac_sh and Aj_sh, the same limitations for the shaded leaf fraction.

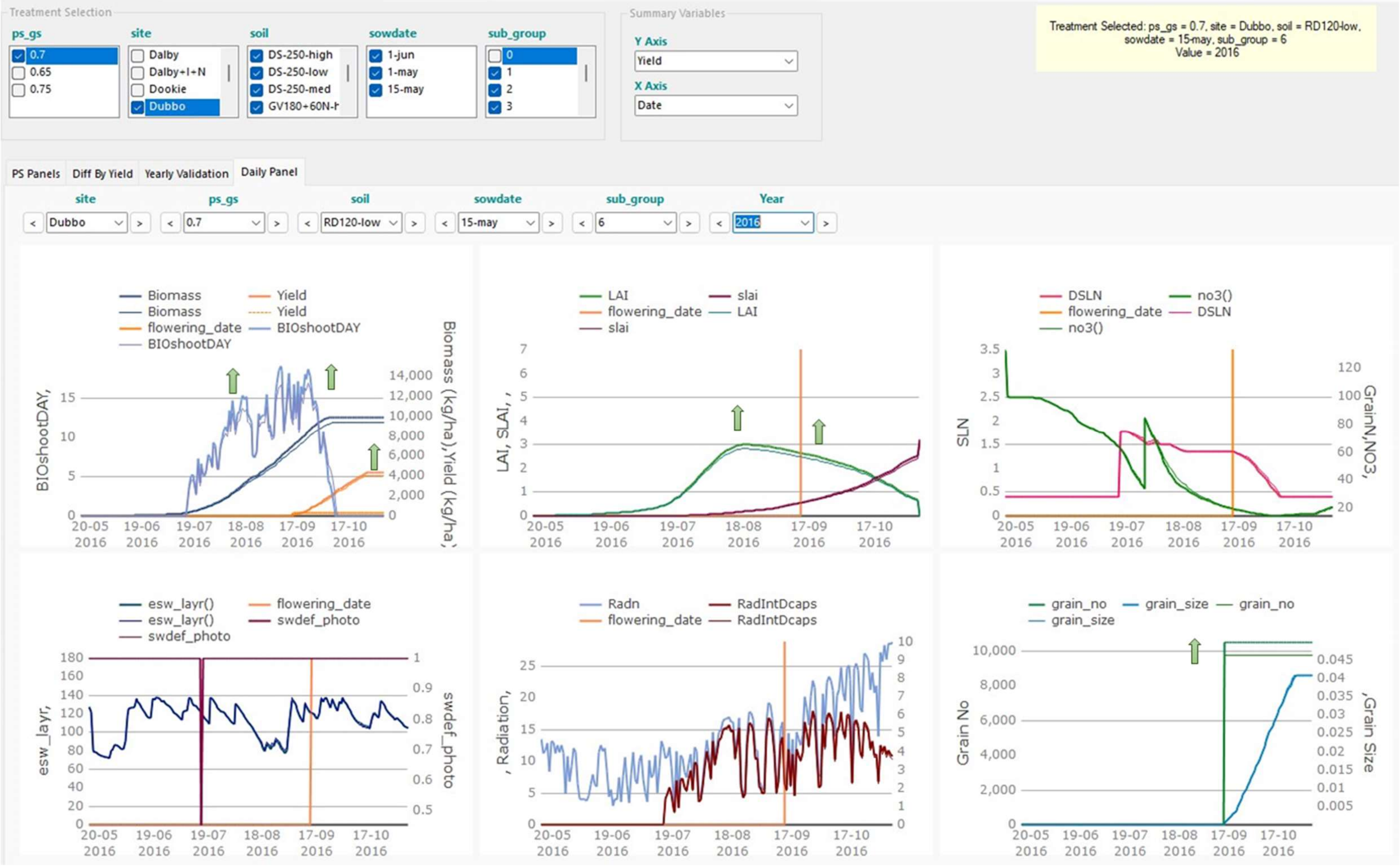


Figure S5. Wheat crop attributes trajectories with abundant water and N. A season selected from the top right-hand corner of the points in Figure 5f of the main text. Baseline trajectories are thin lines; full CCM installation (target 4.2) are thick lines. Arrows highlight changes in trajectories.

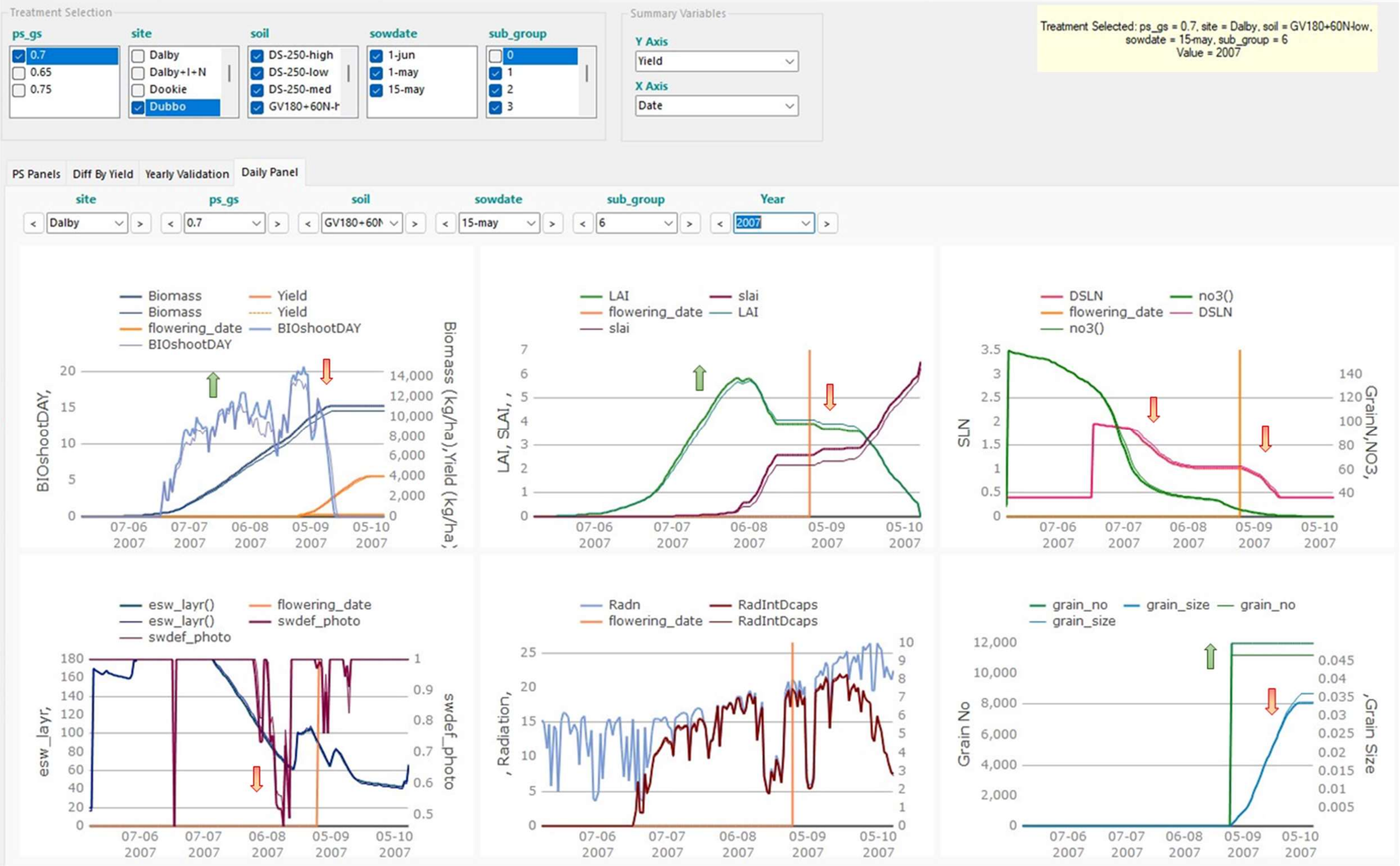


Figure S6. Wheat crop attributes trajectories with exacerbated water stress. A season selected from the bottom right-hand corner of the points in Figure 5f of the main text. Baseline trajectories are thin lines; full CCM installation (target 4.2) are thick lines. Arrows highlight changes in trajectories.

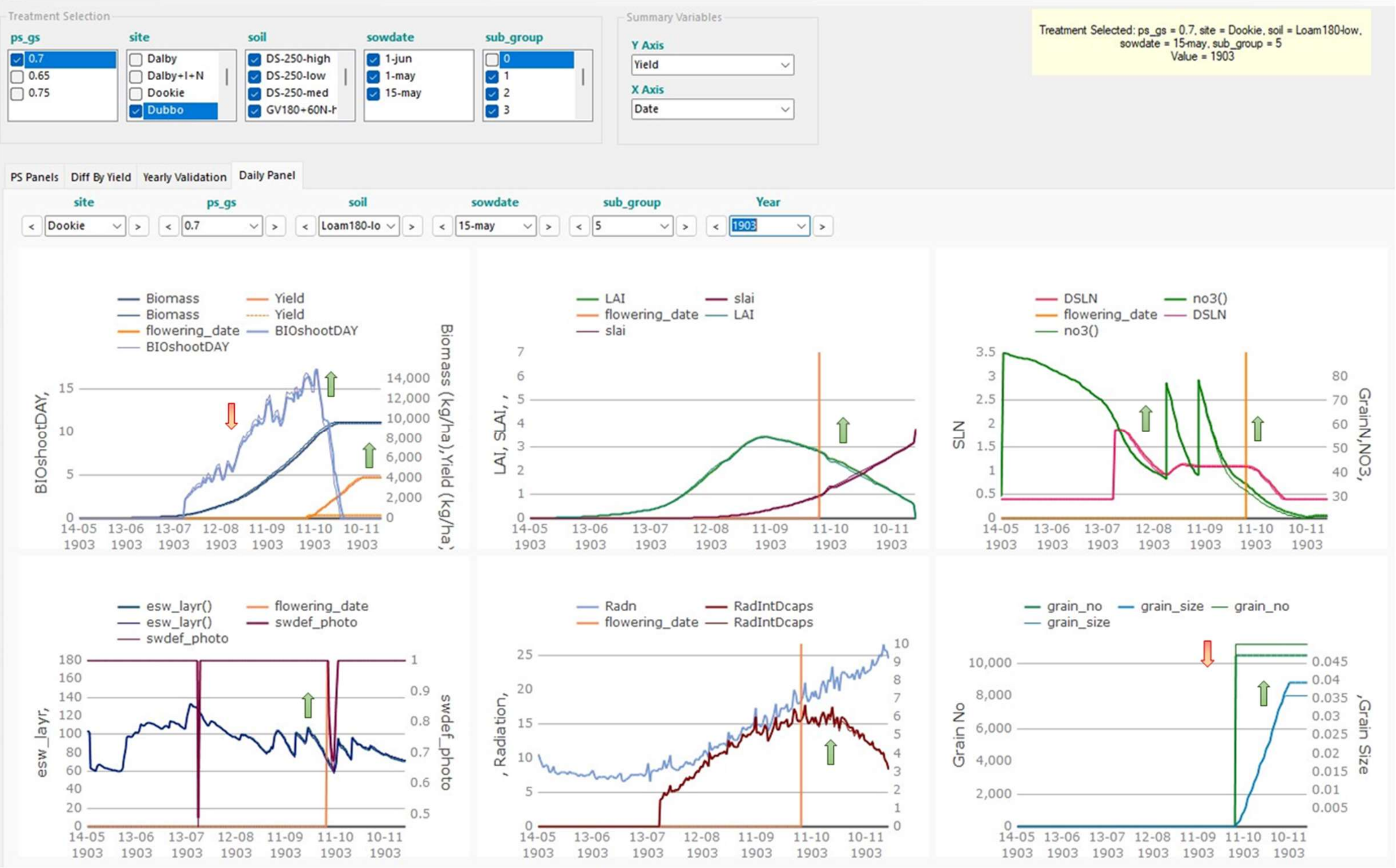


Figure S7. Wheat crop attributes trajectories with reduced growth. A season selected from the top right-hand corner of the points in Figure 5e of the main text. Baseline trajectories are thin lines; HCO3 transporter installation (target 4.1) are thick lines. Arrows highlight changes in trajectories.

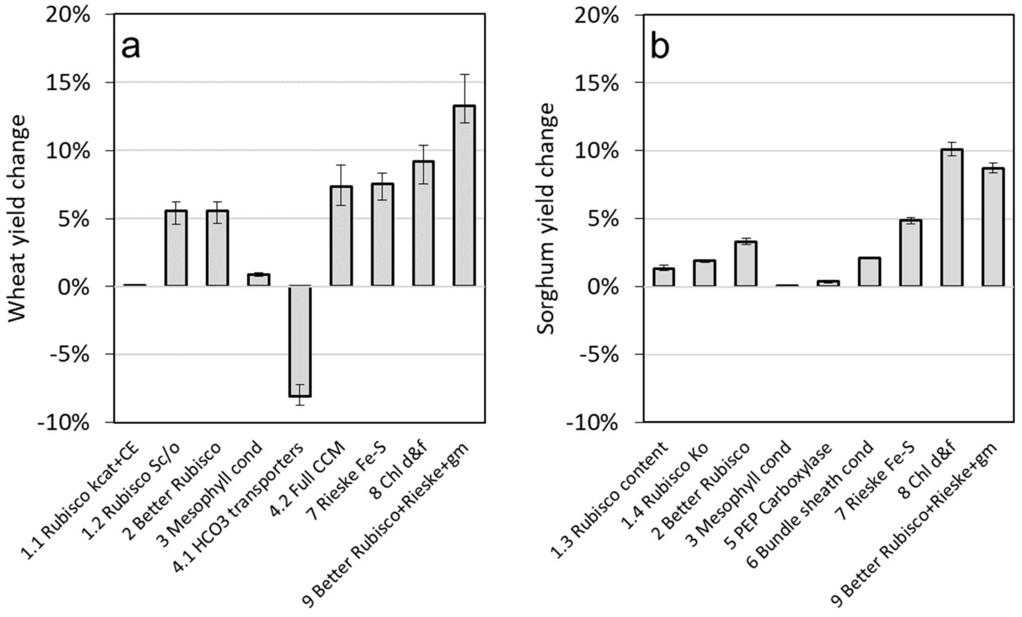


Figure S8. Simulated wheat and sorghum yield change with favourable production conditions. Crops were grown in Dalby with full irrigation and abundant N fertilisation (+I+N) (300kg/ha total). The median sowing date and starting soil water were used (Table S4). Simulated yield data from 120 seasons were used for each leaf photosynthetic manipulation (Table 1). Median values are given by bars. Whiskers show the first and third quartile values.

162

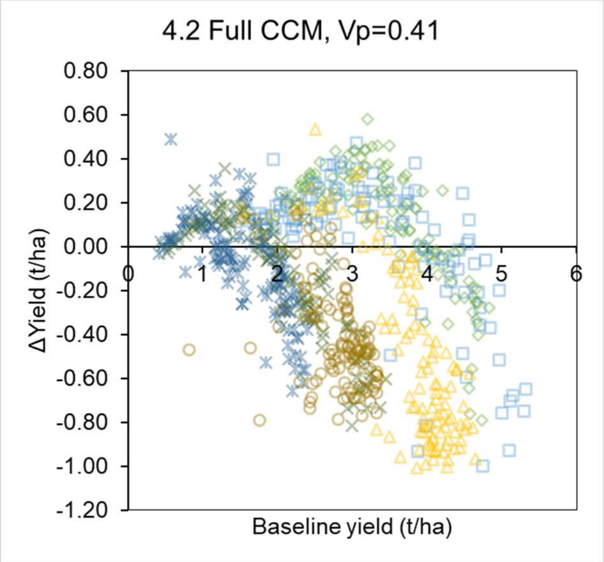


Figure S9. Analogous to Figure 5e in the main text, but with PsiVp=0.41, giving $V_{bmax}$ of 45 instead of the $C_4$ sorghum level at 120. Symbols are different sites (see Figure 5).

163

Table S4. Characteristics of crop production sites, cultivars, and agronomic practices used in simulations.

| Crop type | Production Region | Representative Production Site | Soil type | Soil depth (cm) | PAWC (mm) | Representative sowing dates (Early/Mid/Late) | Representative PAW (mm) at sowing (Low/Mid/High) | Standard crop configuration | Locally adapted nitrogen application (kg N $ha^{-1}$)* |
|---|---|---|---|---|---|---|---|---|---|
| Wheat (Janz: Medium-maturing) | Queensland, northern New South Wales | Dalby (59) | Grey vertosol | 180 | 206 | 1 May/ 15 May/ 1 June | 172/206/206 | 0.25-meter row spacing and 100 plants $m^{-2}$ | 30-130-0-0 |
| | Central, northern New South Wales | Dubbo (48) | Red dermosol | 120 | 142 | | 104/142/142 | 0.25-meter row spacing and 100 plants $m^{-2}$ | 50-50-50[d]-0 |
| | Coastal shires in Victoria | Dookie (39) | Loam | 180 | 109 | | 64/109/109 | 0.25-meter row spacing and 150 plants $m^{-2}$ | 50-40-40[d]-40[c] |
| | South Australia, Victoria Southern New South Wales | Walpeup (32) | Loamy sand | 130 | 134 | | 31/59/113 | 0.25-meter row spacing and 100 plants $m^{-2}$ | 50-20-30[b]-30[c] |
| | North-east inland Western Australia | Merredin (14) | Shallow loamy duplex | 120 | 101 | | 16/37/74 | 0.25-meter row spacing and 100 plants $m^{-2}$ | 30-20-20-30[a] |
| | South-west costal Western Australia | Katanning (8) | Deep sandy duplex | 250 | 74 | | 46/74/74 | 0.25-meter row spacing and 150 plants $m^{-2}$ | 45-20-30-30[a] |
| Sorghum (Hybrid MR-Buster: Medium-maturing) | Central Queensland | Emerald | Black earth | 120 | 150 | 15 Nov/15 Dec/15 Jan | 86/98/109 | 1-meter row spacing and 5 plants $m^{-2}$ | 30-120-0-0 |
| | Southern Queensland | Dalby | Vertosol | 180 | 324 | 15 Oct/15 Nov/15 Dec | 137/200/262 | | 30-120-0-0 |
| | Northern New South Wales (upper) | Moree | Grey clay | 150 | 263 | | 121/168/214 | | 30-120-0-0 |
| | Northern New South Wales (lower) | Gunnedah | | | | | | | 50-100-0-0 |

*Wheat data taken from Chenu *et al.* (2013): initial and applied nitrogen (N) is indicated by '*x*-*y*-*z*-*a*': *x*, antecedent soil N (the baseline N before fertilisation) evenly present throughout soil layers at sowing; *y*, N applied at sowing at 50 mm deep as nitrate, but captured by the soil module of the simulation by adding *y* into the top layer to simplify simulation files; *z* and *a*, N applied as nitrate at the stages 'beginning of stem elongation' and 'mid-stem elongation', respectively. Sorghum data: *x* taken from Hammer *et al.* (2014); *y* calculated by recommended N application given targeting grain yield in the top quartile levels.

[a]If soil PAW > 60 mm at the stage 'mid-stem elongation".

[c]If soil PAW > 60% of PAWC at the stage 'mid-stem elongation'.

[b]If > 80 mm of rainfall from sowing to the stage 'beginning of stem elongation'.

[d]If > 100 mm of rainfall from sowing to the stage 'beginning of stem elongation'.

Table S5. Photosynthesis model parameters and their baseline values used in the $C_3$, single-cell CCM, and $C_4$ photosynthesis models. Values were adapted from Wu et al (2019) with some recalculated using new data. Values in brackets indicate level of the photosynthetic parameter in health, fully expanded leaves with a specific leaf nitrogen (SLN) of 2 g N $m^{-2}$.

| Parameter | Units | $C_3$ wheat | Single-cell CCM wheat: adding bicarbonate transporters | Single-cell CCM wheat: full CCM | $C_4$ sorghum |
|---|---|---|---|---|---|
| $\chi_{Vc}$ ($V_{cmax25}$) | mmol $CO_2$ $mol^{-1}$ N $s^{-1}$ (μmol $m^{-2}$ $s^{-1}$) | 1.45 (158) | 1.45 (158) | | 0.28 (31) |
| $\chi_{Vp}$ ($V_{pmax25}$) | | - | 0.41 (45)* | 1.1 (120)* | 1.1 (120) |
| $\chi_J$ ($J_{max25}$) | | 2.4 (271) | 2.4 (271) | | 2.5 (277) |
| $K_c$ | μbar | 268 | 268 | 2656 | 1194 |
| $K_o$ | μbar | 164697 | 164697 | 136698 | 291378 |
| $S_{c/o}$ | μbar $μbar^{-1}$ | 2799 | 2799 | 1713 | 1339 |
| $K_p$ | μbar | - | 60** | | 74 |
| $\chi_{gm}$ ($g_{m25}$) | mol $CO_2$ $mol^{-1}$ N $s^{-1}$ $bar^{-1}$ (mol $m^{-2}$ $s^{-1}$ $bar^{-1}$) | 0.005 (0.55) | 0.011 ($1.20^{+}$) | | 0.011 (1.20) |
| $g_{bs}$ | mol $m^{-2}$ $s^{-1}$ $bar^{-1}$ | - | ($1^{++}$) | ($0.003^{++}$) | (0.003) |
| $\varphi$ | - | - | 0.75 | | 2 |
| $\alpha_{PSII}$ (single leaf) | - | 0.36 | 0.36 | | 0.26 |
| $\theta$ | - | 0.7 | 0.7 | | 0.68 |
| $C_i/C_a$ | - | 0.7 | 0.7 | | 0.4 |

* In the single-cell cyanobacterial CCM model, this value is the slope ($\chi_{Vb}$) of the linear relationship between the maximum rate of bicarbonate transporters ($V_{bmax25}$) and SLN for SLN greater than a leaf structural N level.

** In the single-cell CCM model, this value is renamed the Michalis-Menten constant of bicarbonate transporter for $CO_2$.

$^{+}$ Conductance for $CO_2$ across the mesophyll cell wall plasmalemma interface

$^{++}$ Conductance for $CO_2$ across the chloroplast envelope.

Table S6. Long-term average cropping area, yield, and total production calculated at the regional level using shire-level data from the Australian Bureau of Statistics. Wheat values are averaged yearly values between 1901 and 2004; sorghum values are averaged yearly values between 1983 and 2015. The area and production values in the AUS Total row are consistent with the Australian Government Department of Agriculture, Water and the Environment (ABSRES) state-level crop data (file access: https://www.agriculture.gov.au/abares/research-topics/agricultural-outlook/data#australian-crop-report-data).

| Crop | Region ID | Region name | Representative Production Site | Area (Mha) | Yield (t/ha) | Production (Mt) | Regional Production Weighting |
|---|---|---|---|---|---|---|---|
| Wheat | 1 | Queensland, northern New South Wales | Dalby | 0.56 | 1.47 | 0.82 | 4% |
| | 2 | Central, northern New South Wales | Dubbo | 2.32 | 1.94 | 4.49 | 24% |
| | 3 | Coastal shires in Victoria | Dookie | 0.16 | 2.62 | 0.43 | 2% |
| | 4 | South Australia, Victoria Southern New South Wales | Walpeup | 2.58 | 1.99 | 5.15 | 27% |
| | 5 | North-east inland Western Australia | Merredin | 2.97 | 1.89 | 5.62 | 30% |
| | 6 | South-west costal Western Australia | Katanning | 1.01 | 2.31 | 2.32 | 12% |
| | AUS Total | | | 9.60 | | 18.82 | 100% |
| Sorghum | 1 | Central Queensland | Emerald | 0.18 | 1.73 | 0.32 | 20% |
| | 2 | Southern Queensland | Dalby | 0.27 | 2.78 | 0.77 | 48% |
| | 3 | Northern New South Wales | Moree, Gunnedah | 0.18 | 2.86 | 0.52 | 33% |
| | AUS Total | | | 0.63 | | 1.61 | 100% |

Table S7. Predicted median percentage change in crop production at the regional level.

| Crop | Region ID | Region name | Representative Production Site | Production (Mt) | 1.1 Rubisco kcat + CE (%) | 1.2 Rubisco Sc/o (%) | 2 Better Rubisco (%) | 3 Mesophyll cond (%) | 4.1 HCO3 transporters (%) | 4.2 Full CCM (%) | 7 Rieske Fe-S (%) | 8 Chl d&f (%) | 9 Better Rubisco + Rieske + gm (%) |
|---|---|---|---|---|---|---|---|---|---|---|---|---|---|
| Wheat | 1 | Queensland, northern New South Wales | Dalby | 0.82 | 1.20 | -0.71 | 0.65 | 0.10 | 3.11 | 4.75 | -1.58 | -1.75 | -0.18 |
| | 2 | Central, northern New South Wales | Dubbo | 4.49 | 1.00 | -0.12 | 1.10 | 0.20 | 1.71 | 5.90 | -0.89 | -0.69 | 0.81 |
| | 3 | Coastal shires in Victoria | Dookie | 0.43 | 0.84 | 0.19 | 1.68 | 0.37 | -0.58 | 3.90 | -0.71 | -0.18 | 2.07 |
| | 4 | South Australia, Victoria Southern New South Wales | Walpeup | 5.15 | 0.89 | 0.08 | 1.30 | 0.22 | 0.84 | 2.87 | -0.42 | -0.14 | 1.63 |
| | 5 | North-east inland Western Australia | Merredin | 5.62 | 0.93 | -0.58 | 0.76 | 0.14 | 1.78 | 1.22 | -1.25 | -0.65 | 0.29 |
| | 6 | South-west costal Western Australia | Katanning | 2.32 | 0.63 | 0.14 | 1.24 | 0.37 | 0.11 | 1.54 | -0.61 | -0.40 | 2.04 |

| Crop | Region ID | Region name | Representative Production Site | Production (Mt) | 1.3 Rubisco content (%) | 1.4 Rubisco Ko (%) | 2 Better Rubisco (%) | 3 Mesophyll cond (%) | 5 PEP Carboxylase (%) | 6 Bundle sheath cond (%) | 7 Rieske Fe-S (%) | 8 Chl d&f (%) | 9 Better Rubisco + Rieske + gm (%) |
|---|---|---|---|---|---|---|---|---|---|---|---|---|---|
| Sorghum | 1 | Central Queensland | Emerald | 0.32 | 0.87 | 0.95 | 1.86 | 0.09 | 0.44 | 1.16 | 2.14 | 3.87 | 4.13 |
| | 2 | Southern Queensland | Dalby | 0.77 | 1.33 | 1.00 | 2.39 | 0.05 | 0.59 | 1.34 | 1.95 | 3.62 | 4.66 |
| | 3 | Northern New South Wales | Moree, Gunnedah | 0.52 | 1.12 | 0.53 | 1.73 | 0.10 | 0.55 | 0.81 | 0.83 | 1.20 | 2.63 |
